# In-silico molecular enrichment and clearance of the human intracranial space

**DOI:** 10.1101/2025.01.30.635680

**Authors:** Marius Causemann, Miroslav Kuchta, Rami Masri, Marie E. Rognes

## Abstract

The mechanisms of intracranial solute transport are fundamental to human brain health, with alterations often linked to disease and functional impairment, and with distinct opportunities for personalized diagnostics and treatment. However, our understanding of these mechanisms and their interplay remains incomplete, in part due to the complexity of integrating insights across scales, between species and from different modalities. Here, we combine mixed-dimensional modelling, multi-modal magnetic resonance images, and high performance computing to construct and explore a high-fidelity in-silico model of human intracranial molecular enrichment. This model predicts the temporo-spatial spreading of a solute within an image-derived geometric representation of the subarachnoid space, ventricular system and brain parenchyma, including networks of surface perivascular spaces (PVSs). Our findings highlight the significant impact of cerebrospinal fluid (CSF) production and intracranial pulsatility on molecular enrichment following intrathecal tracer injection. We demonstrate that low-frequency vasomotion induces moderate CSF flow in surface PVS networks which substantially enhances tracer enrichment, and that impaired enrichment is a direct natural consequence of enlarged PVSs. This openly available technology platform thus provides an opportunity for integrating separate observations on diffusion in neuropil, vascular dynamics, intracranial pulsatility, CSF production, and efflux, and for exploring drug delivery and clearance in the human brain.

## Introduction

The mechanisms underlying molecular transport within the intracranial space are fundamental to human brain health and function. Neurodegenerative diseases such as Alzheimer’s, Parkinson’s, and Huntington’s disease are all associated with abnormal accumulation of protein aggregates together with alterations in transport and clearance characteristics^1–4^. Moreover, sleep and conversely sleep-deprivation play a definite yet enigmatic role in modulating molecular enrichment and clearance^5–9^. In the last decade, established theories have been challenged by new findings revealing a greater degree of molecular movement and exchange^10–14^, as well as substantial variability in enrichment and clearance between individuals and between patient cohorts^7,15–17^. These observations provide distinct opportunities (and challenges) for personalized medicine e.g. for tailored intrathecal delivery of chemotherapy^18^ and for early diagnostics of impaired brain clearance^19,20^. In spite of their importance, our understanding of these mechanisms is incomplete with open challenges and significant debate – in part relating to the translation of knowledge between scales, species, experimental protocols, clinical cohorts, and individuals.

Perivascular pathways along the brain surface and within the brain parenchyma have long been hypothesized to serve a designated role in this context^10,21–27^. Recently, Eide and Ringstad^28^ and Yamamoto et al^29^ demonstrated that perivascular spaces (PVSs) define preferential pathways for molecular transport in humans, with delayed periarterial enrichment in dementia subtypes^28^. Perivascular flow of cerebrospinal fluid (CSF) clearly contributes to this transport, and is inherently associated with vascular pulsations^30–35^. Fluid mechanics considerations point at intracranial pressure differences and shorter wavelength vascular wall pulsations as drivers of directional net flow and convection in the PVS^36–41^, while longer waves such as the pulse wave primarily contribute to oscillatory flow and dispersion^42–47^. In the bigger picture, CSF is produced by the choroid plexus^48–50^, pulsates through the ventricular system, cisterns, and subarachnoid space (SAS) in synchrony with cardiac, respiratory, and neural waves^16,51–58^, and drains via the dural sinuses, meningeal lymphatics, cranial nerves, or other efflux pathways^13^. However, how these physiological factors and physical mechanisms integrate to enhance or impair human intracranial molecular transport over larger spatial scales and longer time scales remain unknown. A related key question is to what extent surface PVSs are separated from the SAS by structural barriers^22,28,59–64^, and in turn to what extent such structural compartmentalization is a prerequisite for effective perivascular transport.

In this study, by leveraging geometric model reduction and mixed-dimensional modelling^65^, structural magnetic resonance (MR) images^66^, and high performance computing, we introduce an integrated computational model of intracranial molecular enrichment and clearance. Focusing on the interplay between perivascular pathways, pulsatility and CSF flow dynamics, the model predicts the temporal evolution and spatial distribution of a solute concentration within a detailed geometric representation of the human SAS and ventricular system, networks of surface PVSs and the brain parenchyma. In terms of transport dynamics, we account for heterogeneous diffusion, dispersive mixing induced by cardiac and respiratory pulsatility in the CSF spaces and PVSs, convective fluid flow driven by CSF production and peristaltic pumping, as well as solute exchange and clearance across semi-permeable membranes. Comparing with glymphatic MRI studies^11,15,67^, the in-silico predictions accurately represent molecular enrichment patterns, timing and intercompartmental distributions. This open platform^68^ thus provides a technological opportunity for qualitatively and quantitatively exploring key open questions relating to molecular movement within the human brain environment such as the role of pulsatility, perivascular pathways, structural compartmentalization, or morphology.

By exploring this high-dimensional parameter space, we propose that the balance between CSF production and intracranial pulsatility is key to shaping the large-scale features of intracranial enrichment patterns, with the potential to span a wide range of individual and cohort variability. Moreover, we predict that CSF production, cardiac- and respiratory pulsatility is not sufficient to explain early perivascular enrichment, but that fluid flow induced by low-frequency vasomotion in surface periarterial spaces (on the order of 10 µm/s) is sufficient, even in the absence of a structural compartmentalization of the PVS. Conversely, enlarged PVSs in the SAS will cause a substantial reduction in cardiac- and vasomotion-driven flow velocities, strongly delay perivascular transport, and thus impair intracranial enrichment. These findings transfer, reconcile, integrate and extend insights from clinical, experimental, and theoretical studies, and lay a new foundation for in-silico studies of personalized intrathecal drug delivery and brain clearance.

## Results

### In-silico predictions of intracranial molecular enrichment and clearance after intrathecal injection

Using previously published multi-modal magnetic resonance imaging (MRI) data^66,69–72^, we construct a multiscale computational representation of the human intracranial compartments consisting of the CSF spaces and brain parenchyma as three-dimensional (3D) domains and with the PVSs surrounding major surface arteries and veins as embedded networks of topologically one-dimensional (1D) curves (Figure 1A–C). We consider a solute concentration field, varying in space and time, in the 3D domains and in the PVS networks, and assume that the solute can cross between these compartments through semi-permeable membranes. As the drivers and modes of intracranial transport are under substantial debate^13,14,73–75^, our first target is to establish a baseline model accounting for a reasonably conservative set of mechanisms and their integrated effect over a timescale of several minutes to a few days. To this end, we assume that the solute will (i) diffuse within all compartments, with diffusivity depending on the effective properties of the relevant medium^76^; (ii) experience significant dispersive effects due to the pulsatile flow of CSF induced by the cardiac and respiratory cycles^43,46,55,77^; and (iii) be convected by a (small) net flow of CSF resulting from production in the choroid plexus with CSF efflux across the upper convexity^78^ and from the peristaltic pumping effect of pulse wave pulsations in surface periarterial spaces^33,40^. Mathematically, this model is represented by a mixed-dimensional system of coupled time-dependent partial differential equations^65^, which we solve numerically with high accuracy using a mass-conserving finite element scheme and the FEniCS finite element software^79,80^ (see Methods). The computational framework and associated software are all openly available^68^.

**Figure 1.**
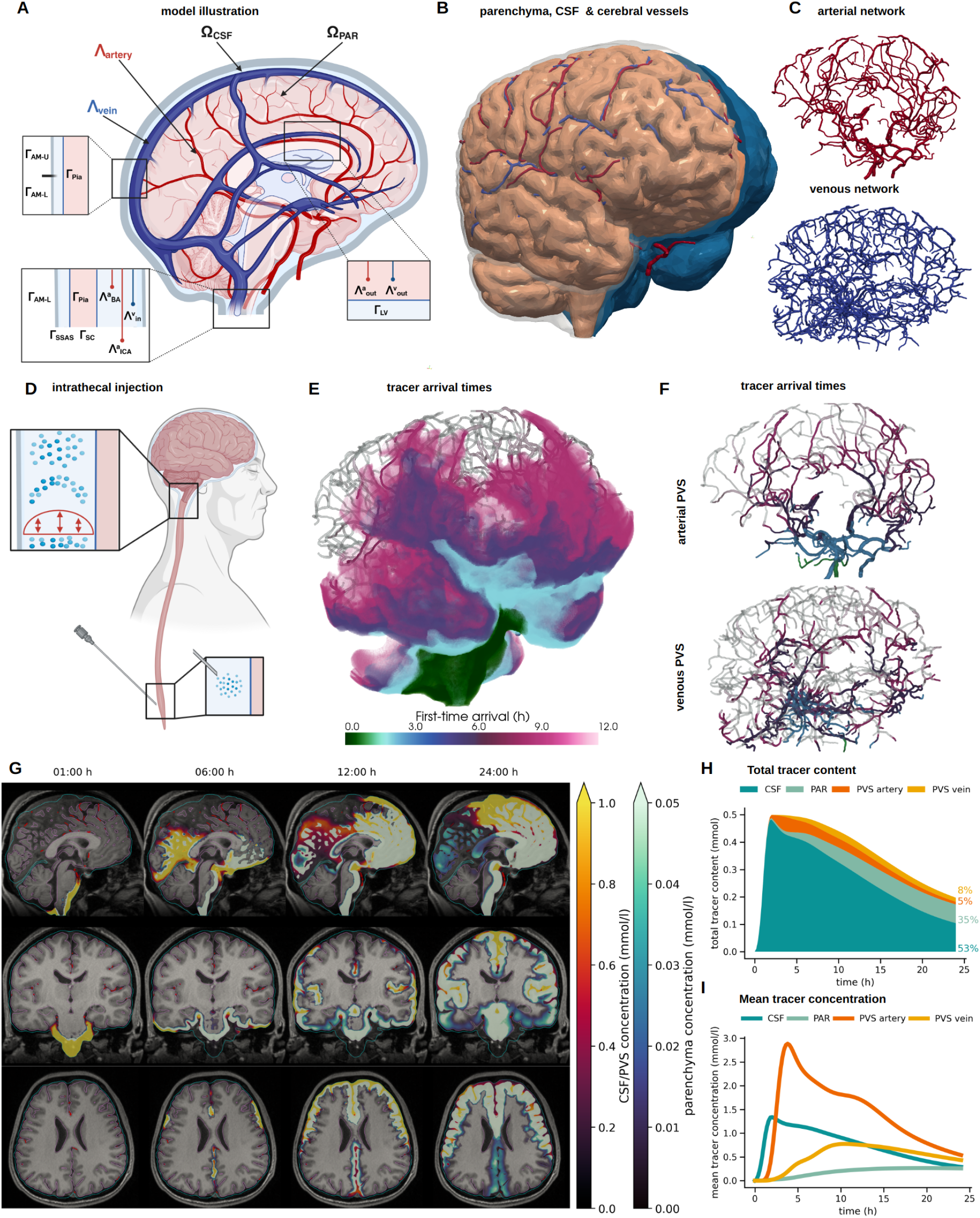
A) Illustration of the spatial domains of the model: CSF-filled space (ventricles and SAS) Ω_CSF_, parenchyma Ω_PAR_ and the PVS surrounding arteries Λ_artery_ and the veins Λ_vein_ and their interfaces and boundaries (see Methods); B) 3D rendering of the actual geometry used in this study parenchyma, CSF (clipped, blue), arterial network (red), venous network (blue)); C) arterial network (red) and venous network (blue); D) illustration of the modelled clinical tracer injection procedure; E) time of simulated first tracer arrival after intrathecal injection (concentration exceeds 0.1 mol/l); F) time of first tracer arrival in the arterial PVS (top) and venous PVS (bottom); G) sagittal, coronal and axial view of computed tracer concentrations over T1-image 1, 6, 12 and 24 h after injection (low concentrations transparent, pial surface in pink, arachnoid membrane in cyan, and arteries in dark red); H) total amount of tracer in each cranial compartment over the first 24 h; I) average tracer concentration in each compartment over the first 24 h.

Simulating a glymphatic MRI protocol^11,15,28^(Figure 1D), we then predict the spreading of 0.5 mmol intrathecally injected Gadobutrol after its appearance at the craniocervical junction (Figure 1E–G). After one hour, the in-silico tracer moves upwards in the SAS frontally of the brainstem, and quickly reaches the supratentorial regions. Here, it spreads both posterior through the quadrigeminal cistern and the longitudinal fissure, and anteriorly through the outer SAS, reaching the top of the cerebral cortex after around 12 hours. After 24 hours, the tracer covers most of the brain surface (with the exception of some posterior regions) and has penetrated substantially into the parenchymal tissue. This pattern is reflected in the mean tracer concentrations in each compartment, where the CSF space reaches its peak of 1.4 mmol/l after 2 hours, followed by the arterial PVS concentration peak (2.9 mmol/l after 4 hours) (Figure 1I). While the mean concentrations in these compartments drop soon after peaking, the parenchymal tissue slowly enriches with tracer over the first 24 hours, with a final value of 0.3 mmol/l. Overall, about 40% of the total amount of tracer remains in the cranium 24 hours post-injection, with the largest share in the CSF (53%), followed by the parenchyma (35%), the venous PVS (8%) and the arterial PVS (5%) (Figure 1H).

### Reduced CSF pulsatility strongly shifts intracranial enrichment patterns

The enrichment patterns observed clinically after intrathecal injection of contrast differ substantially between subjects and between pathological conditions^7,15–17^. These neurological conditions are also associated with alterations in the pulsatile flow of CSF in the ventricular system and SAS^16^. Clearly, key physiological factors such as cardiac pulsatility and respiration easily differ between individuals and between both pathological and physiological states. We therefore next asked to what extent – and how – variations in CSF pulsatility would affect the intracranial enrichment characteristics.

For the CSF dynamics in our baseline model, we combine the net contribution to CSF flow induced by CSF production with the integrated dispersive effects of cardiac and respiratory pulsatile CSF flow (Figure 2A–C). Imposing a constant CSF production of 400 ml/day across the surface of the lateral ventricles, while allowing for efflux across the upper convexity, yields a total CSF pressure drop of 26 mPa (0.00020 mmHg) (Figure 2D) with a maximum flow velocity of 1.85 mm/s (in the aqueduct) and a mean velocity in the SAS of 2.6µm/s (Figure 2E). To examine the dispersive effects of pulsatile CSF flow, we employ the theory of shear-augmented dispersion together with computational fluid dynamics to determine a dispersion factor

**Figure 2.**
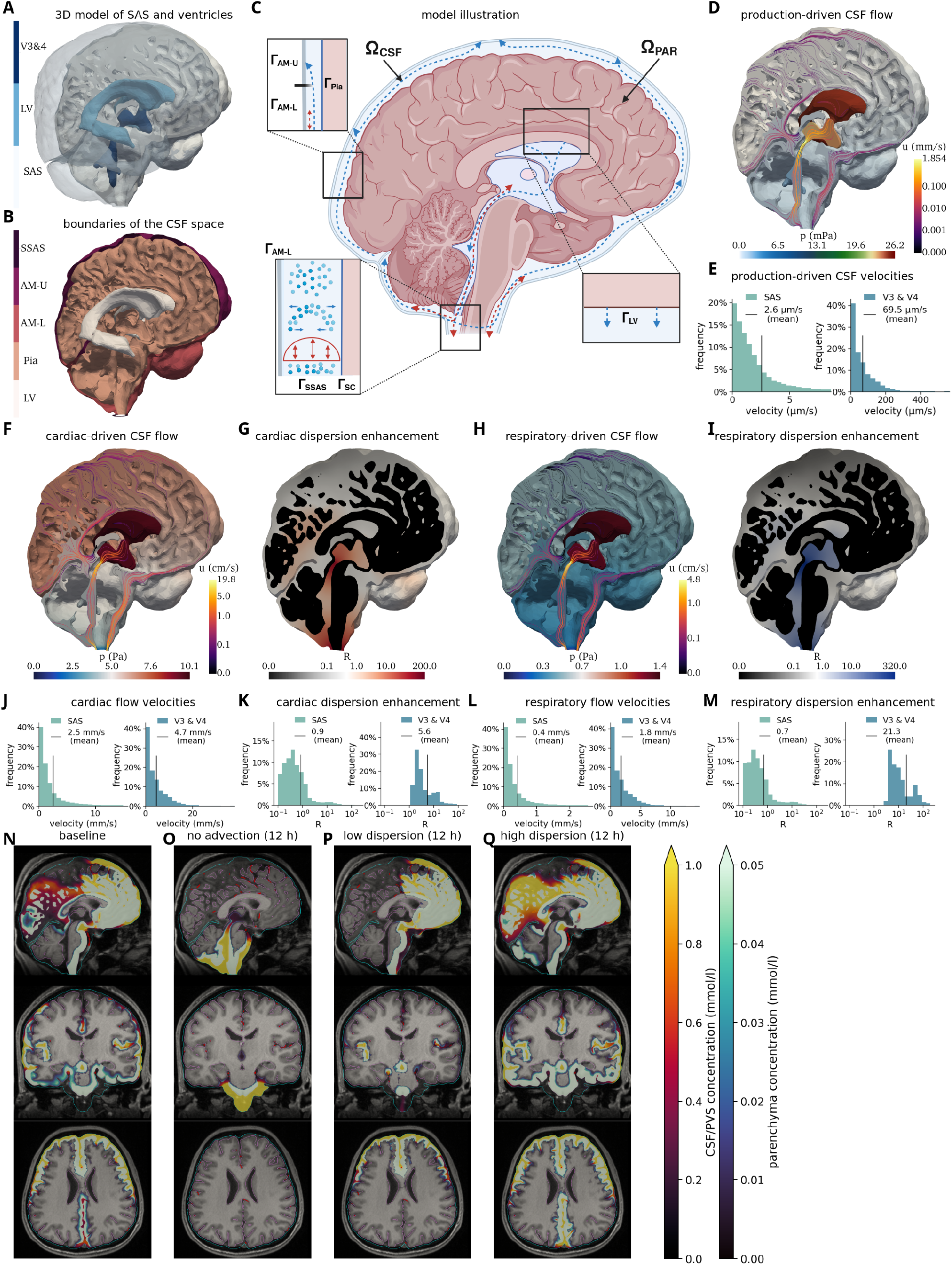
The balance between CSF pulsatility and production shapes molecular enrichment patterns. A) CSF spaces: the third and fourth ventricle (V3&4), lateral ventricles (LV) and SAS; B) Boundaries and interfaces: the spinal subarachnoid space (SSAS), upper and lower arachnoid membrane (AM-U and AM-L), pia (Pia) and lateral ventricles (LV); C) Schematic of the CSF flow model; D) CSF flow induced by CSF production with outflow allowed across the upper convexity (AM-U); E) Histograms of the production-induced CSF velocities in the SAS and third and fourth ventricle (V3 & V4) in terms of the relative frequency of the mean flow speed in each computational cell weighted by its volume; F) CSF flow induced at peak systolic blood inflow; G) Cardiac-induced dispersion factor *R*_*c*_; H) CSF flow induced at peak respiratory expansion; I) Respiration-induced dispersion factor *R*_*r*_, *D* = (1 + *R*_*c*_ + *R*_*r*_)*D*^Gad^; J–M): Histograms of the cardiac-induced flow velocities (J), cardiac-induced dispersion factors (K), respiratory flow velocities (L), respiratory dispersion factor (M); N–Q): sagittal, coronal and axial planes of tracer concentration after 12h for different model variations: baseline (N), no CSF production (O), low dispersion (P), and high dispersion (Q).

*R* enhancing the effective solute diffusivity (see Methods). For the cardiac contribution, we compute CSF pressure and flow fields at peak systolic blood inflow, corresponding to a reduction in CSF space volume at a total rate of 6 ml/s^56,81^ in the SAS and 0.31 ml/s across the lateral ventricle surface^55^. This scenario sets up a pressure drop of 10 Pa (0.075 mmHg) between the lateral ventricles and the spinal SAS and a maximum flow velocity of 19.8 cm/s (Figure 2F, J). Assuming a cardiac frequency of 1 Hz, we infer that this cardiac-induced pulsatile CSF flow increases the effective diffusion by more than two orders of magnitude in the aqueduct and near the cisterna magna, but has little effect (*R* < 1) in most of the SAS (Figure 2G, K). For the respiratory contribution, we employ the same methodology, but with a total rate of 1 ml/s^82^ in the SAS and 0.121 ml/s^83^ in the lateral ventricles, yielding a respiratory peak flow volume of 1.121 ml/s at the craniocervical junction. While the resulting flow velocities are only about one fourth of their cardiac-induced counterparts (Figure 2H, L), respiratory dispersive mixing reaches a factor of up to 320 due the lower respiratory frequency of 0.25 Hz (Figure 2I, M).

Now, to examine how CSF pulsatility affects the enrichment patterns, we consider three variations of the baseline: (i) no CSF production, (ii) reduced pulsatility and thus decreased dispersion (0.1 × *R*), and (iii) higher pulsatility with increased dispersion (10× *R*). Without CSF production, transport is considerably delayed (Figure 2N, O). Even after 12 hours, tracer remains in the subtentorial regions around the cerebellum, the brain stem and in the surrounding CSF. Interestingly, the lack of CSF production instead allows tracer to travel upwards through the ventricular system, reaching the third ventricle after around 12 hours. On the other hand, if the CSF pulsatility is reduced, we observe rapid transport towards the upper convexity of the cranium, as in the baseline model, but the tracer spreads exclusively within the anterior regions (Figure 2P). This feature can be attributed to the CSF flow bifurcation posterior to the ambient cistern (Figure 2D): without sufficient diffusion, the tracer is unable to cross into the posterior SAS. Indeed, with higher dispersion, the tracer moves through the quadrigeminal cistern and further upwards into the longitudinal fissure, with also enrichment of the cerebellum (Figure 2Q).

### Perivascular flow shapes and accelerates molecular enrichment

The PVSs are recognized across species as critical pathways for solute transport in and around the brain, and thus as potential targets for enhancing brain drug delivery and metabolic waste clearance. However, whether CSF flows more rapidly in PVSs and what the forces and mechanisms required to drive such flow are, remain as key points of debate^14,27^. Motivated by experimental observations in animal models^10,31–33^, there is now a remarkable body of literature on modelling perivascular fluid flow and transport^36–47^. Here, we ask how these proposed mechanisms would translate from idealized geometries to human vascular networks and moreover evaluate their integrated effect in the context of intracranial solute transport.

Our periarterial network extends from the internal carotid arteries and basilar artery through up to 18 bifurcations to reach upstream network ends located within the SAS or up to 6 mm inside the parenchyma (Figure 1A–C, Figure 3A) and includes major surface arteries of radius 0.5–1.4 mm. Imposing the pressure field induced by CSF production at the periarterial network ends induces slow steady CSF flow of variable direction in these PVSs, with an average velocity of 0.08 µm/s (antegrade) and a maximum velocity of 0.55 µm/s (Figure 3B). On the other hand, traveling waves of arterial wall motion (Figure 3C), such as the pulse wave or other vasomotion^9,84–87^, also induce net directional flow in the PVS^39–41,45,88^ – of magnitude and direction depending on the amplitude, frequency, and length of the waves and the characteristics of the perivascular network^40^. Applying a semi-analytic model of the net flow induced by peristalsis in perivascular networks^40^ (see Methods), we estimate that the cardiac pulse wave alone, traveling at a frequency of 1 Hz with a wavelength of 2.0 m and a 1% wall displacement^89^, will induce mainly antegrade PVS flow with an average net velocity of 0.92 µm/s while reaching up to 7.31 µm/s near larger, ventral vessels such as the MCA (Figure 3D). The same theory predicts that strong ultraslow vasomotions, if traveling antegrade at 0.1Hz with a wavelength of 0.02 m and a 10% displacement^87^, will induce both retrograde and antegrade net PVS flow with an average velocity of 13.05 µm/s and maximum velocity 54.44 µm/s (Figure 3E).

**Figure 3.**
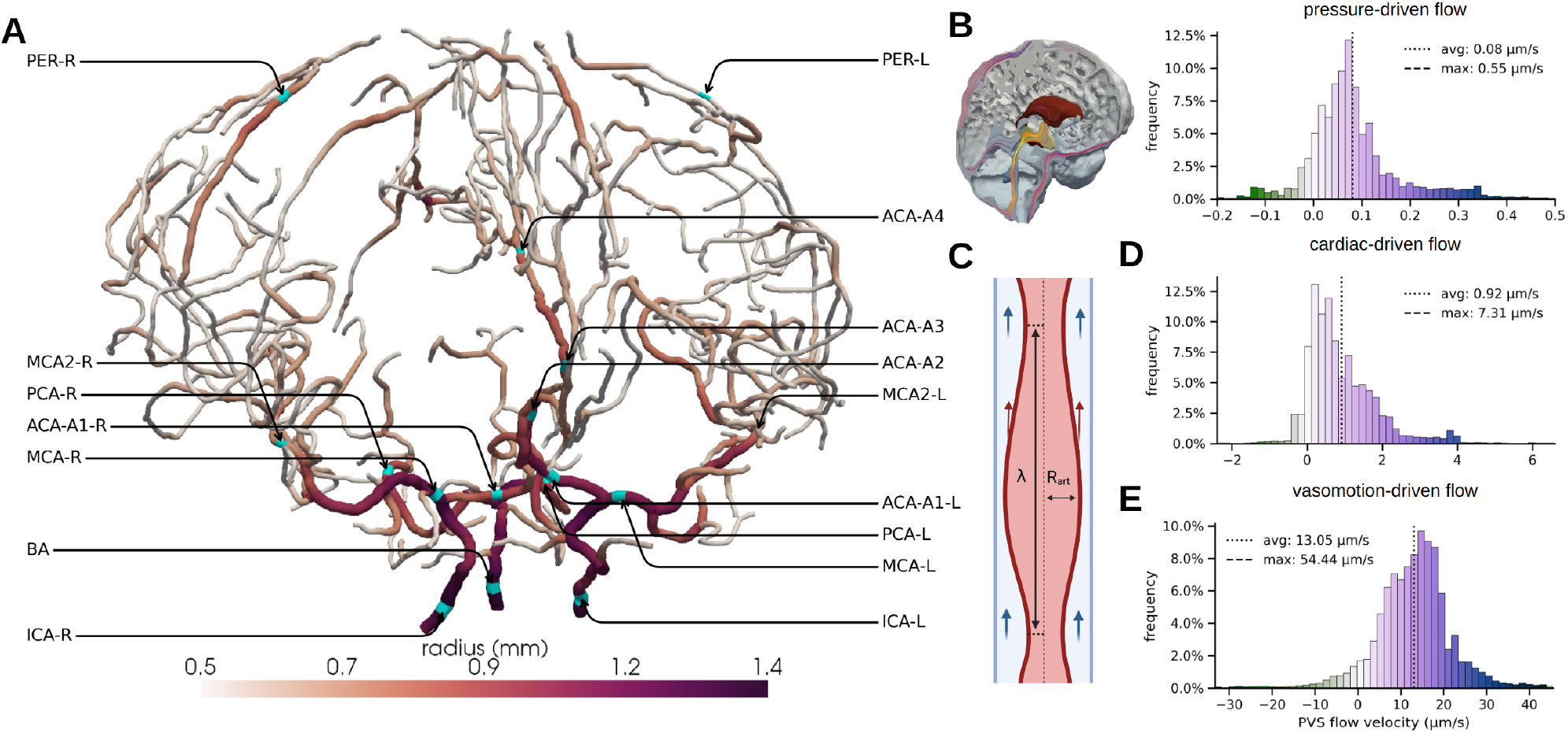
A static pressure gradient and peristaltic pumping are potential drivers of perivascular flow. A) Surface arterial network colored by arterial radius with vessel locations labeled; B) Net PVS flow induced by CSF production (cf. Figure 2D) – histograms show the relative frequencies of local periarterial velocities. Positive values indicate antegrade PVS flow, while negative values indicate retrograde PVS flow; C) Illustration of the concept of peristaltic pumping; D) Estimated net PVS flow induced by pulse wave peristaltic pumping; E) Estimated net PVS flow induced by vasomotion/slow wave peristaltic pumping.

We study the effect of such rapid PVS flow on intracranial molecular enrichment by comparing the more conservative baseline model and a high PVS flow model, where the former still includes the net flow contributions from CSF production and the cardiac pulse wave, while the latter additionally includes the ultraslow vasomotion contribution. In both models, tracers were first observed at the basal artery after 48 minutes (Figure 4A), defined by their first-time arrival (FTA), the time at which the concentration first exceeds 0.1 mmol/l. At all upstream locations along the middle cerebral arteries (MCAs) and anterior cerebral artery (ACA), we found substantially reduced FTAs with higher PVS flow (Figure 4B), up to 2.5 hours earlier in the left M2 segment of the MCA. PVS concentrations peaked before the surrounding tissue (0:48h vs 1:12h at the MCA and 1:00h vs 1:48h at MCA2) with high PVS flow, while these peaks occurred nearly simultaneously in the baseline model. The earlier appearance and the delay in time-to-peak between the PVS and surrounding tissue clearly indicates the directionality of tracer enrichment – it first arrives in the PVS and subsequently spreads into the tissue and SAS (Figure 4C, D), and especially so with higher PVS flow. These observations also hold on an aggregated level: computing the total amount of tracer in the PVS as a function of the distance to the arterial network roots, we find accelerated tracer transport with higher PVS flow, especially at earlier time points (2–9 hours) (Figure 4E). Moreover, differences in tracer transport along the PVS translate to altered enrichment patterns on the whole-organ scale. For instance, after 4-6 hours, the faster-moving tracer in the PVS is clearly visible in the space adjacent to the PVS of the ACA, MCA, and other arteries (Figure 4G). In contrast, there are no clear signs of early enrichment surrounding the PVS in the baseline model (Figure 4F).

**Figure 4.**
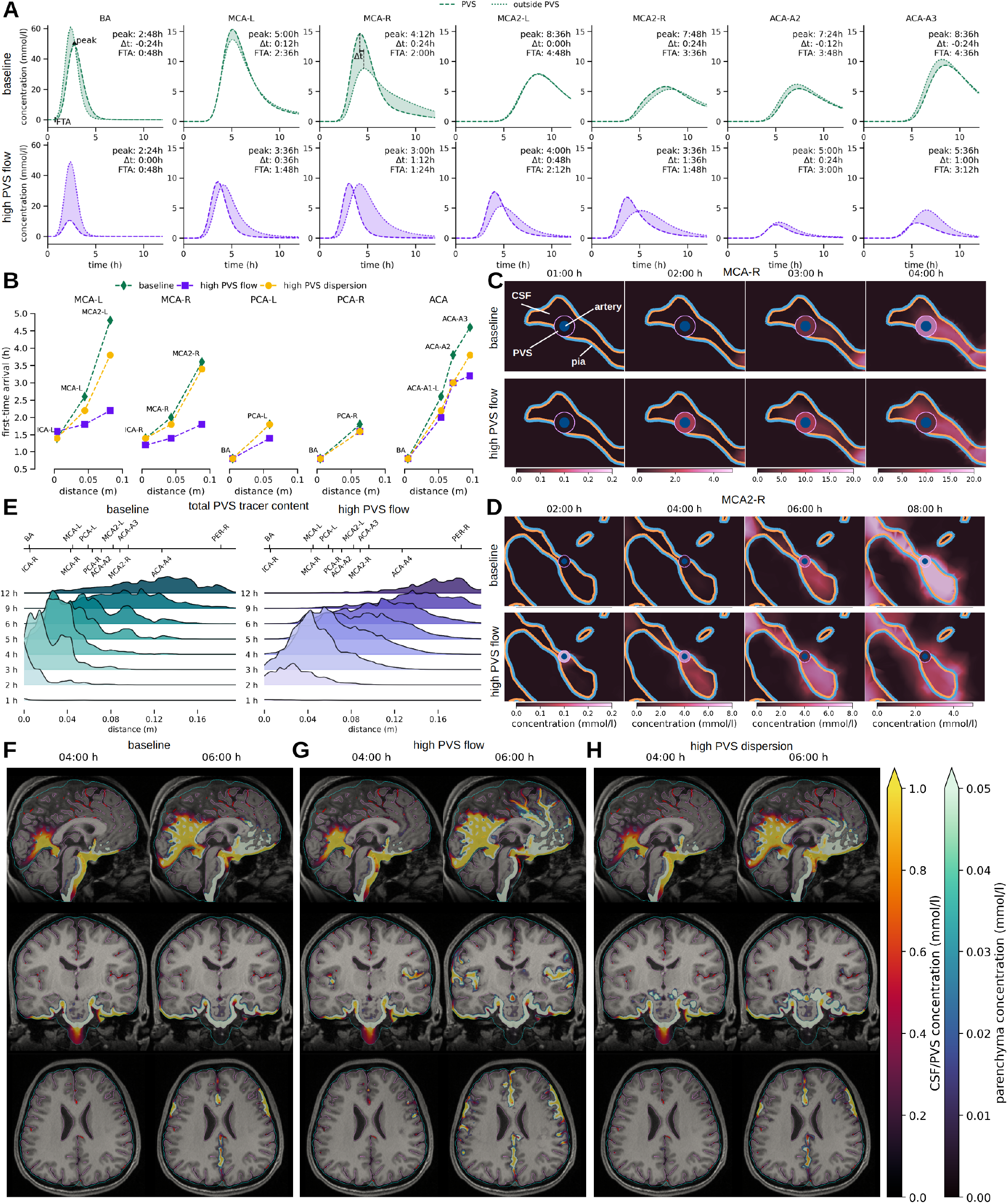
Perivascular flow shapes and accelerates molecular enrichment. A) Comparison of mean tracer concentration in the PVS and over the outer PVS surface at key locations of the arterial tree, for the baseline (lower) and high PVS flow (upper) models, with labels: “peak”: time-to-peak (h), “Δ*t*”: time difference between peaks (h), and “FTA”: first-time tracer arrival (h). B) first-time arrival for the main trunks of the periarterial network over the distance from the closest network root node (BA, ICA-R or ICA-L for the baseline, the high PVS flow and the high PVS dispersion models); C, D) 2D slices showing zoom-in on the region surrounding the MCA-R and MCA2-R for the baseline (upper) and high PVS flow (lower) models. Innermost circle indicates the artery (blue), the surrounding annulus is the PVS (color represents concentration value), the surroundings are parenchyma and/or CSF spaces, separated by the pia (blue: towards the parenchyma, pink: towards the CSF)); E) Total amount of tracer in the periarterial spaces as a function of the distance to the nearest network root over time for the baseline (left) and the high PVS flow (right) models; F–H) sagittal (upper) and coronal (lower) views of the simulated tracer concentrations overlayed on the T1-weighted MR image after 4 and 6h for the baseline (K), high PVS flow (L) and high PVS dispersion (M) models (concentration opacity linearly increasing with concentration value, pial surface outlined in pink, arachnoid membrane in cyan, and arteries in dark red). An interactive visualization of the baseline and the high PVS flow model can be found here, and here.

A natural question is whether early PVS enrichment could be the result of increased dispersion rather than net flow in the PVS^42,42,43,46,86^. Interestingly, even increasing the dispersion factor by 100× in the PVS, induced only minor changes in the global spreading rate compared to the baseline (Figure 4F, H), thus indicating a negative answer to this question.

### Structural versus functional compartmentalization of perivascular spaces

Human and rodent observations indicate that tracers concentrate in perivascular spaces surrounding the pial and subarachnoid vasculature^22,28,32,33,60,90^. However, it remains unclear whether such enrichment patterns necessitate a structural barrier, such as a membrane with limited permeability, or if the patterns could result from enhanced flow or mixing in these areas alone. We therefore next investigate how the permeability of the interface between the PVS and the surrounding CSF affects tracer enrichment around the major arterial trunks (Figure 5A–B). To this end, we compare models with high and low permeabilities, representing a highly permeable PVS-CSF interface (functional compartmentalization) and a less permeable interface (structural compartmentalization), respectively. For the low permeability model, we set the permeability to 3.8 · 10^−7^ m/s, consistent with previous estimates for the endfoot sheath surrounding penetrating arterioles^91^, which we consider to be a lower bound for the surface PVS-CSF permeability. For the high permeability model, we increase this permeability by a factor of 100. For both scenarios, we consider the high PVS flow regime as examined in the previous section.

**Figure 5.**
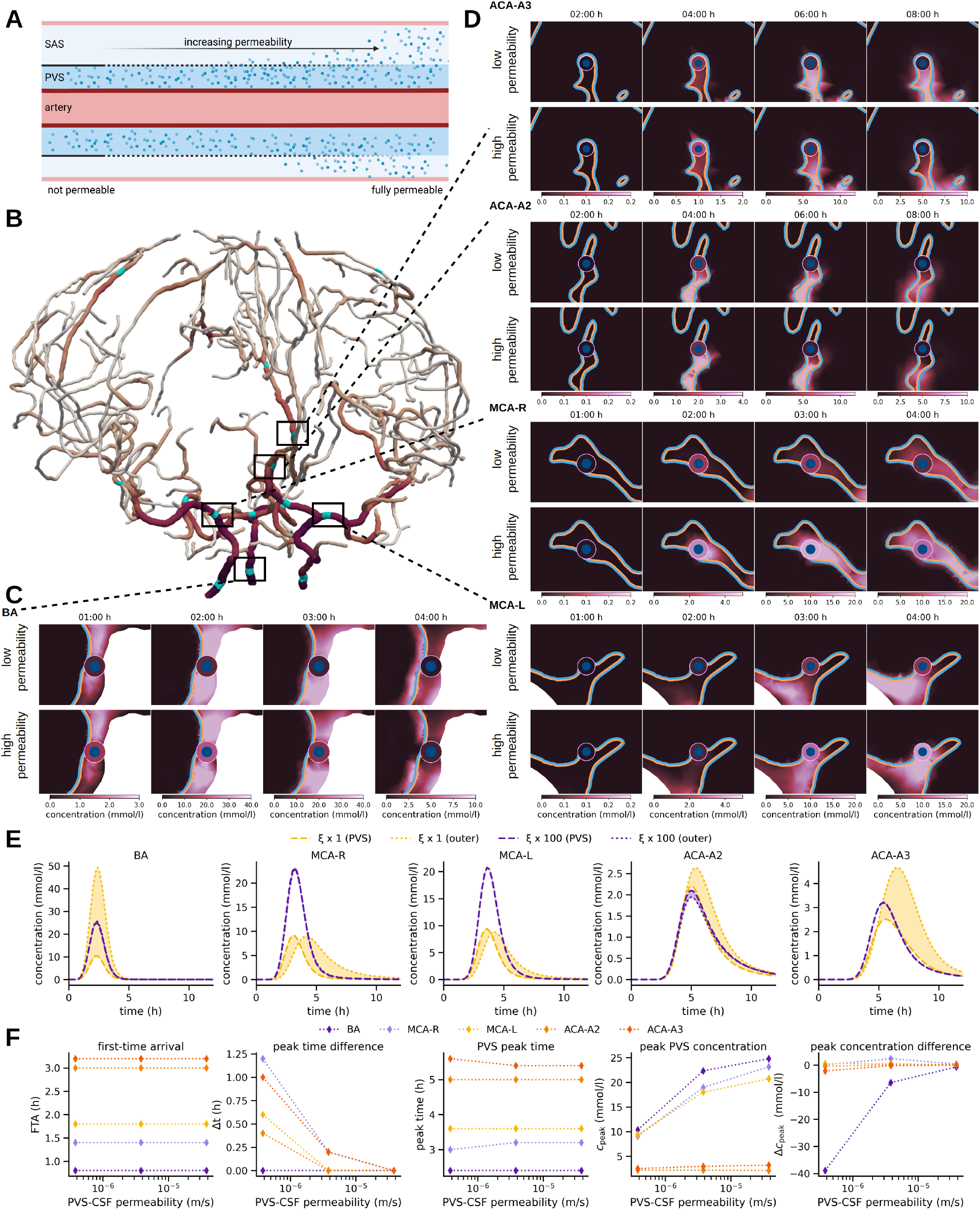
Does early perivascular enrichment rely on a structural compartmentalization of the PVS? A) The PVS-CSF membrane permeability regulates the exchange of solutes between the PVS and surrounding CSF spaces; B) The arterial network with highlighted regions of interest; C) 2D slices showing zoom-in to the region surrounding the basal artery after 1, 2, 3, and 4 hours for a low (above) and high (below) PVS-CSF permeability model. Innermost circle indicates the artery (blue), the surrounding annulus is the PVS (color represents concentration value), the surroundings are parenchyma and/or CSF spaces, separated by the pia (blue: towards the parenchyma, pink: towards the CSF)); D) As in (C) for regions surrounding (from upper to lower) the ACA-A3, ACA-A2, MCA-R and MCA-L segments; E) Comparison of mean tracer concentration in the PVS (dashed) and over the outer PVS surface (dotted) for the BA, MCA-R, MCA-L, ACA-2, and ACA3 segments over time for the high (purple) and low permeability (yellow) models; F) Effect of varying the PVS-CSF permeability on first-time-of-arrival (FTA), time difference (Δ*t*) between concentration peaks in the PVS and surrounding tissue, time of PVS peak concentration, PVS peak concentration and difference in peak concentration between PVS and surrounding tissue.

At the basal artery (BA), tracer appears within one hour in the surrounding CSF in both models (Figure 5C, E). With a more permeable PVS-CSF interface, tracer quickly crosses from the CSF into the PVS, resulting in similar peak concentrations of 25 mmol/l in the PVS and at its outer surface, while values up to 50 mmol/l are attained in the vicinity. In contrast, the low permeability model exhibits substantially lower perivascular tracer enrichment, with a maximum concentration of 10 mmol/l after approximately two hours. For the middle cerebral arteries (MCA-R and MCA-L), tracer appears first in the PVS in both models (Figure 5D, E), though with higher concentrations in the high permeability model (up to 24 mmol/l). We also observe that the higher permeability allows the tracer to leak out of the PVS and spread within the Sylvian fissure to a greater extent. With a small delay, more tracer appears via the CSF pathway (Figure 5D). At the anterior cerebral artery (ACA-A2), the first tracer arrival in the PVS occurs after 3 hours, with a peak at 5 hours. The concentration peak in the PVS is tailed by a higher peak in the CSF (Figure 5E), indicating tracer arrival through a second pathway; i.e., directly through the CSF-filled space outside of the PVS. Similar patterns are observed for ACA-A3.

Summarizing and quantifying these observations (Figure 5F), we find that neither the time of first arrival nor the time-to-peak are substantially affected by the PVS-CSF permeability at any of the periarterial segments considered (BA, MCAs, ACAs). However, the concentration difference between the PVS and its surrounding CSF quickly decreases with higher permeability, as does the time lag between the concentration peaks in these domains. Thus, fast transport along the PVS is not contingent upon a structural compartmentalization of the PVS, whereas a sharp concentration gradient between PVS and CSF is unlikely without a restricting barrier.

### Enlarged PVSs delay periarterial and intracranial molecular enrichment

Enlarged PVSs are associated with impaired periarterial tracer transport and more generally with cognitive decline as well as certain subtypes of dementia^28,92^. A key question is thus whether enlarged PVSs alone leads to delayed perivascular and intracranial tracer enrichment. To address this question from a fluid and transport dynamics perspective, we ask our in-silico model to predict the integrated effect of perivascular dilation on PVS flow velocities and tracer enrichment patterns (Figure 6A). Again we compute CSF production-, cardiac-, and vasomotion-induced net PVS flow velocities, but now in a perivascular geometry with an increased outer radius 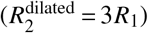 compared to the previous 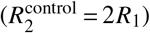, which corresponds to a 2.67× increase in PVS cross-section area. The enlargement yields notable and non-uniform changes in the flow dynamics. On the one hand, the pressure-induced flow velocities increase from a mean of 0.08 µm/s to 0.33 µm/s due to the reduced effective resistance of the PVS. However, the peristaltic pumping becomes considerably less effective: the cardiac and vasomotion-induced net CSF flow velocities drop from a mean of 0.92 µm/s to 0.17 µm/s and from 13.05 µm/s to 2.33 µm/s, respectively (Figure 6B–D). In total, the net CSF velocity reduces, which in turn alters the tracer enrichment within the dilated PVS. Notably, we observe later tracer arrival in all MCA segments, with up to 96 min later arrival in the MCA2s for the high PVS flow scenario (Figure 6E, F). For the baseline model, which omits the vasomotion-induced net PVS flow, the effect is less pronounced in the MCA but clearly persists and is evident in the MCA2s (Figure 6F). Finally, we note that the larger volume of the dilated PVS results in a greater total accumulation of tracer in the PVS from 3–24 hours and reduced enrichment of the parenchyma at 6 and 12 h, while the mean PVS concentration is initially lower, but later exceeds the baseline model (Figure 6G,H).

**Figure 6.**
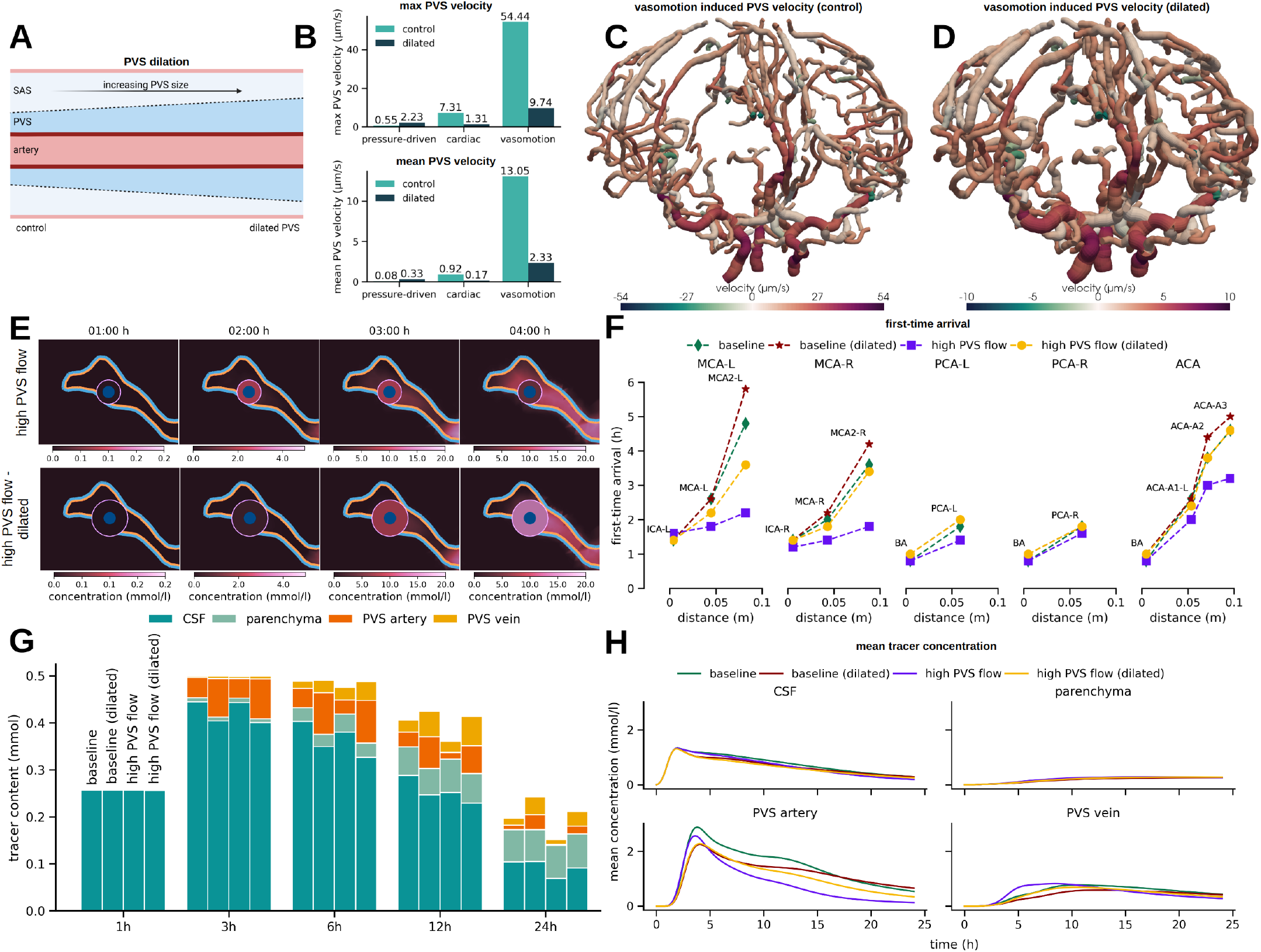
PVS enlargement reduces net PVS flow and delays tracer transport. A) Modelling dilated PVS by extending the outer PVS boundaries; B) mean and max PVS flow velocities with normal/control and dilated PVS; C–D) 3D rendering of vasomotion-induced PVS flow in the normal (C) and dilated (D) periarterial networks; E) 2D slices showing zoom-in to the region surrounding the MCA-R after 1, 2, 3, and 4 h with normal (lower) and dilated (upper) PVS. Innermost circle indicates the artery (blue), the surrounding annulus is the PVS (color represents concentration value), the surroundings are parenchyma and/or CSF spaces, separated by the pia (blue: towards the parenchyma, pink: towards the CSF)); F) First-time arrival for different branches of the arterial tree for the baseline model (with pressure-driven and cardiac-induced net PVS flow) and the high PVS flow model (also including vasomotion-induced net PVS flow) for normal and dilated PVS; G) total tracer content in the CSF, parenchyma and arterial and venous PVS at 1,3,6,12 and 24 h for the four models from F; H) mean concentration in the CSF, parenchyma and arterial and venous PVS over time for the four models from F.

## Discussion

We have presented a high-fidelity in-silico model of molecular enrichment and clearance in human intracranial spaces, enabling tailored predictions of the influence of CSF space and (peri)vascular morphology, physiological factors such as cardiac and respiratory pulsatility, as well as of pathological conditions such as enlarged PVSs. A key observation is that the balance between CSF production and intracranial pulsatility significantly shapes the large-scale features of molecular enrichment, with e.g. ventricular tracer reflux after intrathecal injection in the absence of CSF production, and preference towards anterior brain regions with reduced pulsatility. Indeed, the simulated distribution patterns cover a substantial range of the individual variations observed clinically^15^. In terms of perivascular pathways, we find that even moderate CSF flow on the order of 0.1–1.0 µm/s in the surface PVSs results in earlier tracer enrichment around major cerebral arteries. This effect is more pronounced – with up to a three-hour difference in time of arrival – with mean PVS flow rates of approximately 10 µm/s. Our models of CSF flow in perivascular networks, based on first principles and asymptotic analysis, predict net flow of such magnitudes and both antegrade and retrograde flow within the perivascular network, occurring with or without pressure differences at the network ends. The PVS may thus function as highways facilitating rapid transport, even in the absence of structural barriers, though a sharp concentration difference between the PVS and surrounding tissues is unlikely without such barriers. Finally, dilated PVS will cause a substantial reduction in cardiac- and vasomotion-driven flow velocities, which in turn impedes perivascular transport.

While molecular enrichment of the brain and surrounding CSF spaces have been extensively studied in animal models, especially in murine models in connection with studies of the glymphatic system, there are fewer reports of contrast enrichment in human subjects over 0–24h. We therefore mainly compare our in-silico predictions with the series of papers by Eide and colleagues^6,11,15,28^ and Watts et al^67^. Ringstad et al.^15^ report of tracer enrichment in a centripetal pattern, primarily in regions near large cerebral surface arteries, peaking in the CSF spaces between 6 and 9 hours, with parenchymal tracer content still increasing after 24 hours, but with large individual differences. Watts et al^67^ report of a similar enrichment pattern and a concentration peak in the SAS of ≈ 0.5 mmol/l (relative to a total amount of 1.0 mmol) occuring after 10 to 15 hours. Our in-silico enrichment patterns agree with these observations, though with an earlier rather than later peak in the CSF spaces (2 hours after first tracer appearance at the craniocervical junction). In particular, we note that the substantial individual variation observed clinically is comparable to the variation in in-silico enrichment patterns associated with reduced CSF production or reduced intracranial pulsatility. We find that about 20% of the tracer reaching the cranium enters the parenchyma, which aligns with the reported peak concentration values of 0.5 mmol/l in the SAS and 0.1 mmol/l in the parenchyma reported by Watts et al^67^. In terms of perivascular tracer enrichment, Eide and Ringstad^28^ report of first-time tracer appearance in the PVS of the MCA after 37.8 ±47.0 (M1), 53.1 ±50.5 (M2) and 82.6± 61.5 (M3) min after injection, thus with just a 15.3 min delay between M2 and M1. Comparing with our FTAs in the MCA segments, we note that the baseline model predicts a more-than 3 hour difference in FTA between the M1 and M2. In contrast, the high PVS flow scenario predicts a 24 min delay between these segments. Moreover, the high PVS flow scenario predicts FTAs of 84–108 min (M1) and 108–132 min (M2) after tracer appearance at the craniocervical junction. Assuming a tracer travel time from the point of injection to the craniocervical junction of 13 min (as reported by Eide and Ringstad^28^), our predictions fall at the upper end of the clinical ranges. This can partly be attributed to the limited caudal extend of our PVS network, with the lowest point located approximately 4 cm above the craniocervical junction due to imaging constrains. We interpret these results as indicative of a net CSF flow in human surface PVS beyond what would be induced by CSF production and cardiac peristalsis alone.

The existence, magnitude, and directionality of flow in surface and parenchymal PVSs have been the subject of active debate for decades^10,14,21,24,30–34,36–41,44,47,84,93^. In mice, Mestre et al^33^ observe CSF flow in surface perivascular regions surrounding branches of the MCA with average flow speeds of 18.7 µm/s and net flow in the antegrade direction, but also regions with retrograde flow. Similarly, Bedussi et al.^32^ report of oscillatory flow, with an average (net) CSF velocity of 17 ± 2 µm/s in the antegrade direction. Using physics-informed neural networks, Boster et al^34^ estimate PVS velocities of 12.75± 6.25 µm/s. Less is known about the magnitude and directionality of CSF flow in surface PVSs in humans. Here, assuming that human surface PVS follow the major surface vessels and are CSF-filled but otherwise open regions of width proportional to the vessel radius, our estimates indicate that the CSF pressure differences due to CSF production, cardiac peristaltic pumping and low-frequency vasomotion all yield spatially-varying net CSF flow with both antegrade and retrograde PVS segments. This observation is thus in agreement with the experimental observations^32,33^, and inherently supports the original notion that antegrade and retrograde solute transport along PVSs may coexist^21^. In terms of magnitude, the contribution from CSF production and cardiac wall motion are small (0.08 µm/s and 0.92 µm/s on average), but intriguingly, our estimates of the contribution from vasomotion (13.05 µm/s) are comparable to the experimental reports. We do note that this estimate is based on the presence of rhythmic waves of vasomotion at a frequency of 0.1Hz, a wavelength of 20 mm, and a wall displacement of 10% – in the antegrade direction, all values currently associated with significant uncertainty^84,87,94^.

To what extent are surface PVSs in communication with the surrounding SAS and to what extent are these structurally separated compartments? Originally, Weller and coauthors^22,59,90^ identified thin sheaths of pial cells surrounding human surface arteries and penetrating arterioles (but not veins or venules). On the other hand, Bakker and coauthors report that the subarachnoid space, the cisterns, ventricles and penetrating periarteriolar spaces form a continuous CSF space^60^. Thorne and colleagues study the architectures of rat perivascular spaces in detail, emphasizing the presence of openings (stomata) on the interface between the vasculature and the CSF within the SAS^26,61^. Further, Mestre et al^62^ study the properties of pial perivascular spaces in mice, and report of pial cells forming sheaths for larger surface arteries and partially cover smaller surface arteries, with higher coverage in ventral SAS regions. Our results demonstrate that the PVSs may act as rapid transport pathways even with only a partial barrier between the PVS and surrounding CSF, thus contributing to quantifying the properties of the human PVS-CSF interface^28^.

Lifestyle factors such as sleep^5,6,8,9,86,95,96^, exercise^97,98^, and alcohol intake^99^ affect molecular enrichment and clearance of the brain. In particular, pial arteries display higher-amplitude low-frequency vasomotion during NREM and intermediate sleep states, while REM sleep is associated with reduced PVS width^86^. Peristaltic pumping is more effective under both of these configurations, resulting in higher estimated net PVS flow velocities^40^. Even higher PVS velocities would amplify the PVS flow effects simulated here, with earlier arrival times along the PVSs, clear enrichment in adjacent spaces, and accelerated molecular enrichment overall. Moreover, sleep is linked with low-frequency oscillations in human CSF flow^54^. As we have shown here, the level of dispersion in the CSF spaces strongly shapes molecular enrichment patterns. While we, at baseline, account for the dispersion induced by cardiac and respiratory pulsatility, the high dispersion scenarios considered can be viewed as representative of additional pulsatility induced by e.g. neural waves^54,57^ or other respiration patterns depending on activity level. More broadly, we consider this as a key strength of the current in-silico framework: allowing for integrating and exploring the mechanistic macroscale implications of changes in pulsatility at smaller spatial or temporal scales.

In terms of limitations, our primary constraint modelling-wise is the geometric representation of the vascular (and therefore also perivascular) networks. Concretely, we account for no vessels below the original MRI resolution limits, and thus only a part of the cerebral surface vasculature and effectively not the cortical vessels. In addition, and again due to lack of resolution, there is uncertainty associated with the location of the vessel segments relative to the CSF spaces and parenchyma, with some vessels surrounded by both domains and some vessels completely entering the parenchyma before returning to the SAS. Higher resolution MRI data (T1w and e.g. time-of-flight, QSM or other modalities) in more subjects would immediately allow for more detailed studies of this aspect. As a result of the lack of an accurate representation of the cortical and subcortical white matter vasculature, we have considered a conservative transport model within the parenchyma, modelling extracellular diffusion alone. To compensate, we have focused on reporting detailed predictions in the CSF spaces and surface PVS after not more than 24 hours. In particular, we note that, we observe tracer accumulation at the perivascular network ends after 12 hours, similar to that previously observed for microspheres^32,33^, which here can be interpreted as an artifact of the incomplete network creating up a barrier for tracer movement. Previously, we have found that enrichment within the parenchyma is well-represented by ≈3× enhanced diffusion augmented by local clearance e.g. across the blood-brain barrier or diffusion-convection with a complex pattern of spatially-varying convective velocities of 1-10 µm/min^95^. We will incorporate such transport mechanisms within the parenchyma in future work. Finally, we also note that we simulate the intracranial distribution of a tracer concentration appearing at the craniocervical junction rather than as an intrathecal injection. We therefore overestimate the amount of tracer entering into the intracranial spaces; however, due to the linearity of the mathematical model, it is valid to directly interpret the in-silico concentrations relative to the total amount of tracer.

In conclusion, our findings transfer insights from experimental studies and theoretical analysis to the in-silico human setting, reconcile seemingly conflicting observations in particular relating to directionality of perivascular flow, and integrate different physical mechanisms across across spatial and temporal scales. The complete simulation pipeline is openly available, including interactive visualization of simulation results for all model variations^68^. Looking ahead, this platform establishes a foundation for in-silico studies of molecular movement within the human brain environment, such as tailored predictions of intrathecal chemotherapy delivery or personalized diagnostics of brain clearance capacity.

## Methods

### Intracranial compartments: ventricular system, SAS, and brain parenchyma

From T1-weighted MR images of a 26-year-old, healthy male volunteer^66^, we first automatically segment the brain parenchyma and CSF spaces, including the ventricular system and SAS, using Synthseg^100,101^ (Figure 1A–B). We next manually adjust the segmentation to accurately represent the connections and barriers between CSF spaces (aqueduct, median aperture, tentorium cerebelli), smoothen, and finally extract surface representations of the outer (arachnoid) boundary, the pial membrane and other interfaces. Conforming to these surface and interface representations, we generate a tetrahedral mesh Ω representing the complete intracranial volume as the union of the parenchyma Ω_PAR_ and CSF spaces Ω_CSF_, using fTetWild^102^ (Figure 2A–B). The resulting computational mesh consists of 233592 vertices and 1290131 mesh cells, which vary between 0.22 mm and 8.9 mm in mesh cell diameter. The parenchyma has a total volume of 1318 ml, whereas the ventricles and the outer SAS contribute 32 ml and 329 ml, respectively, to a total CSF volume of 361ml, in agreement with recent estimates^103^.

### Periarterial and perivenous spaces

To represent surface networks of periarterial and perivenous spaces, we use Kiminaro^104^ to separately skeletonize time-of-flight angiography (ToF) and quantitative susceptibility mapping (QSM) images from the same human subject^66^. This technique yields networks of one-dimensional curves, each curve Λ^*i*^ indicating the centerline of a blood vessel segment and labeled with its lumen radius 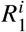 (Figure 1C, Figure 3A). We also assign an outer radius 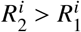 to each segment *i*, form the annular cylinder ensheathing Λ^*i*^ and define the union of these as the PVS. The periarterial network graph consists of 12708 edges connected at 12562 inner nodes and with 147 end nodes, and the associated domain is denoted by Λ_*a*_. We identify three of the end nodes – corresponding to the two internal carotid arteries (ICAs) and the basilar artery (BA) – as root nodes, and designate the other ends as leaf nodes. The resulting perivenous network graph consists of 24829 edges connected at 23881 inner nodes and with 949 end nodes, and the associated domain is denoted by Λ_*v*_. Finally, we label the outer surface of the PVSs associated with Λ_*a*_ and Λ_*v*_, by Γ_*a*_ and Γ_*v*_ respectively.

### Molecular transport equations

We model diffusion, convection, and exchange of a molecular solute in the PVS networks Λ_*a*_ and Λ_*v*_, in the CSF spaces Ω_CSF_ and in the brain parenchyma Ω_PAR_ via a mixed-dimensional transport model^65,105^ over a timescale of minutes to hours (Figure 1A–C, Figure 2A–C). Specifically, for *t* > 0, we solve for a concentration *c* = *c*(*x, t*) in the 3D intracranial compartments (*x ∈* Ω_CSF_, *x ∈* Ω_PAR_) and for a cross-section averaged concentration 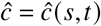 in the periarterial and perivenous networks (*s ∈* Λ_*a*_, *s ∈* Λ_*v*_) such that the following equations hold:

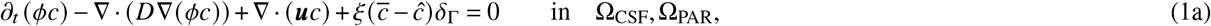

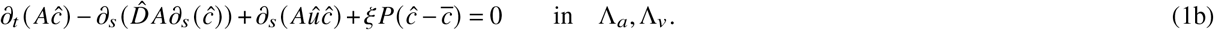

In (1), 0 < *ϕ* ⩽ 1 is the fluid volume fraction, *D* and 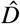 are effective diffusion coefficients, and *u* and 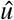 are fluid (CSF/ISF) velocities, all defined in the respective compartments; *P* = 2*πR*_2_ is the perimeter and 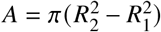 the area of the PVS cross-sections, *δ*_Γ_ is a Dirac term concentrated on the interfaces Γ_*a*_, Γ_*v*_, the overlined *c* denotes the average of *c* over cross-sections of Γ_*a*_, Γ_*v*_, and *ξ* is a permeability allowing for transfer/exchange over the interfaces Γ_*a*_, Γ_*v*_ between the periarterial and perivenous networks and their surroundings. Moreover, we model the membrane between the parenchyma and the CSF spaces as a semi-permeable membrane with permeability coefficient *β*_pia_. For a complete description of this mathematical model and simulation scenarios, see Appendix A.1. The initial concentrations in the three-dimensional domain Ω and networks Λ_*a*_, Λ_*v*_ are all set to zero.

### Intracranial influx after intrathecal injection

To represent molecular influx into the intracranial compartments following intrathecal injection, without modelling the spinal compartment explicitly, we prescribe a time-dependent molecular influx over the interface towards the spinal CSF compartment with the condition

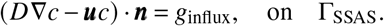

Here we set

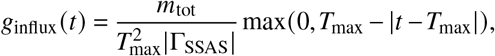

which expresses a hat-shaped influx function with a peak at *T*_max_ and a total tracer injection of *m*_tot_ (Table 1).

**Table 1.**
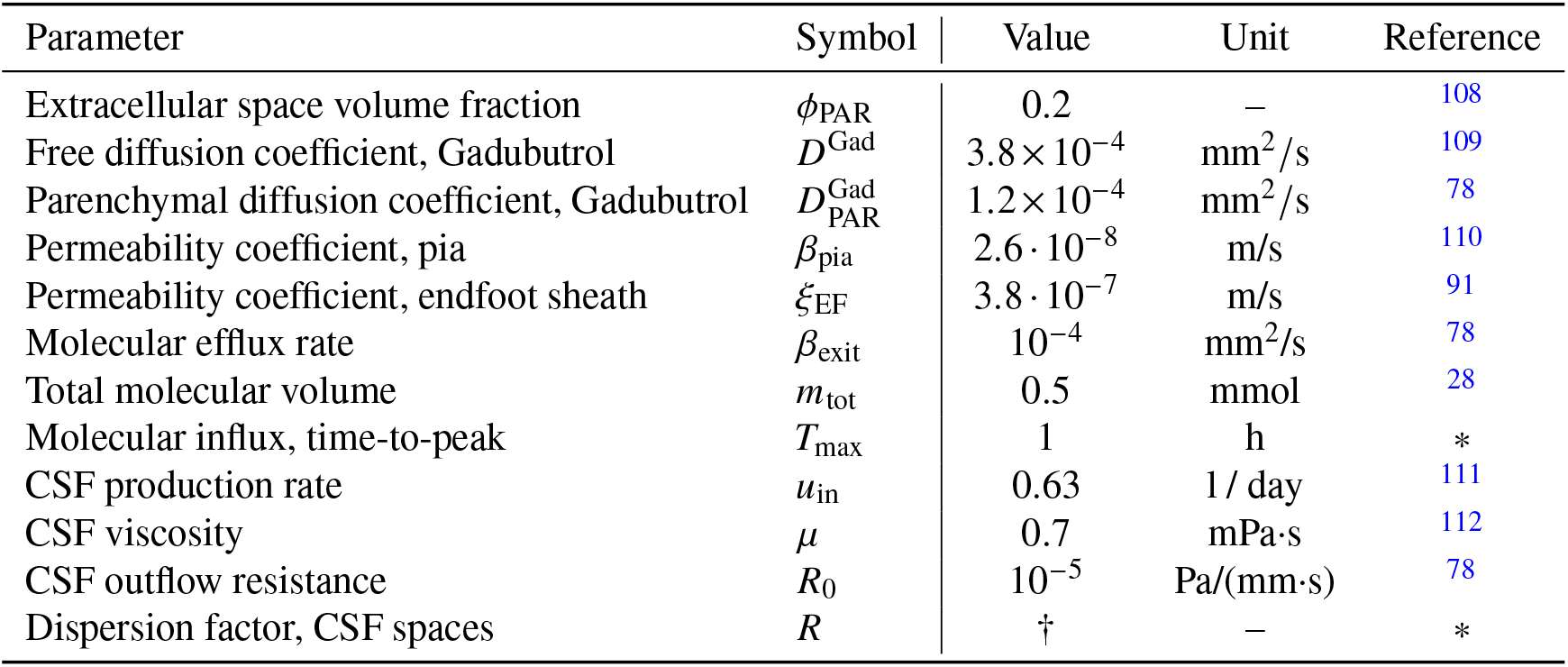
Model parameters. †/* denotes computed/estimated within this work.

| Parameter | Symbol | Value | Unit | Reference |
| --- | --- | --- | --- | --- |
| Extracellular space volume fraction | $\phi_{\text{PAR}}$ | 0.2 | – | 108 |
| Free diffusion coefficient, Gadubutrol | $D^{\text{Gad}}$ | $3.8 \times 10^{-4}$ | $\text{mm}^2/\text{s}$ | 109 |
| Parenchymal diffusion coefficient, Gadubutrol | $D_{\text{PAR}}^{\text{Gad}}$ | $1.2 \times 10^{-4}$ | $\text{mm}^2/\text{s}$ | 78 |
| Permeability coefficient, pia | $\beta_{\text{pia}}$ | $2.6 \cdot 10^{-8}$ | $\text{m/s}$ | 110 |
| Permeability coefficient, endfoot sheath | $\xi_{\text{EF}}$ | $3.8 \cdot 10^{-7}$ | $\text{m/s}$ | 91 |
| Molecular efflux rate | $\beta_{\text{exit}}$ | $10^{-4}$ | $\text{mm}^2/\text{s}$ | 78 |
| Total molecular volume | $m_{\text{tot}}$ | 0.5 | $\text{mmol}$ | 28 |
| Molecular influx, time-to-peak | $T_{\text{max}}$ | 1 | $\text{h}$ | * |
| CSF production rate | $u_{\text{in}}$ | 0.63 | $1/\text{day}$ | 111 |
| CSF viscosity | $\mu$ | 0.7 | $\text{mPa}\cdot\text{s}$ | 112 |
| CSF outflow resistance | $R_0$ | $10^{-5}$ | $\text{Pa}/(\text{mm}\cdot\text{s})$ | 78 |
| Dispersion factor, CSF spaces | $R$ | † | – | * |

### Molecular clearance from the intracranial space and perivascular network ends

We model molecular clearance, into the meningeal lymphatics or other pathways, across the upper, outer (arachnoid) boundary (Figure 1A, Figure 2C), proportional to the concentration in the SAS with a rate constant *β*_exit_. We set a no-flux condition at the end nodes of the perivascular networks, not allowing for direct molecular clearance there.

### CSF flow in the SAS and ventricular system

We model CSF as an incompressible Newtonian fluid at low Reynolds and Womersley numbers via the Stokes equations with viscosity *µ*, and account for the convective contribution of the flow induced by CSF production in the choroid plexus. Specifically, we compute the velocity *u*_CSF_ and associated pressure *p*_CSF_ in the SAS and ventricular system induced by a constant production at a rate of *u*_in_ across the lateral ventricle walls and with efflux across the upper, outer (arachnoid) boundary with efflux resistance *R*_0_; and then set *u* = *u*_CSF_ in Ω_CSF_ in (1) (Table 1, Figure 2A–E, see also Appendix A.1). We do not model bulk flow within the brain parenchyma, except within the perivascular network segments extending below the pial surface, and thus set *u* = 0 in Ω_PAR_ in (1).

### Dispersion in the CSF spaces

The pulsatile flow of CSF in the SAS and ventricular system, associated with the cardiac and respiratory cycles, substantially enhances molecular diffusion through dispersion^42,43,46,77,106,107^. To account for these dispersive effects, while bridging from the pulsatile flow time scale of seconds to a molecular transport time scale of hours, we compute cardiac and respiratory dispersion factors *R*_*c*_ and *R*_*r*_, both spatially-varying, in Ω_CSF_ (Figure 2A) by combining computational estimates of the respective peak CSF pressure gradients with theoretical estimates for shear-augmented (Taylor) dispersion^43,106,107^ (see Appendix A.3 for a complete description). Combining these effects, we then set *R* = *R*_*c*_ + *R*_*r*_ and define the effective diffusion coefficient *D* = (1 + *R*) *D*^Gad^ in Ω_CSF_ in (1) in the baseline model.

### Perivascular fluid flow induced by CSF pressure differences

To estimate the contribution also to perivascular fluid flow from CSF production, we impose the fluid pressure *p*_CSF_ induced by CSF production, computed in the SAS and ventricular system, at the end nodes of the periarterial network located within the SAS. For the network end nodes located within the parenchyma, we compute a harmonic extension of the CSF pressure field (by solving a Laplace equation in Ω_PAR_ with *p*_CSF_ as the boundary value), and set the corresponding value at these end nodes. We then numerically solve a system of hydraulic network equations to compute 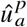 defined over Λ_*a*_; i.e., the pressure-difference induced CSF flow velocity in the periarterial network (see also Appendix A.1.4). We repeat this procedure to compute a corresponding velocity 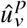 in the perivenous network Λ_*v*_.

### Perivascular fluid flow induced by arterial wall motion

Using the theoretical framework introduced by Gjerde et al.^40^, we also compute analytic estimates for the time-average perivascular flow rate ⟨*Q*^′^⟩ induced by peristalsis in the arterial network Λ_*a*_; i.e. the net flow induced by traveling waves of arterial wall motion of frequency *f*, amplitude *ε* and wave lengths *λ* (Figure 3C). The corresponding contribution to the periarterial flow velocity is defined for each segment Λ_*i*_ as 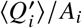 where 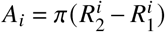 is the cross-section area of the PVS segment and 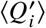 is the estimated mean flow rate (see A.4 for more details). Two wall motion patterns are considered, yielding 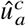 and 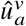 respectively: *cardiac* pulsations with *f* = 1.0Hz, *λ* = 2 m and *ε* = 1% (Figure 3D), and very low-frequency *vasomotion* with *f* = 0.1Hz, *λ* = 0.02m, and *ε* = 10% (Figure 3E). No flow induced by peristalsis is considered for the perivenous network. In the baseline model, we combine the contributions from CSF production and cardiac peristalsis to set 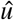 in (1) as 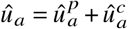 in Λ_*a*_, and 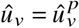 in Λ_*v*_.

### Model and material parameters

Model and material parameters are summarized in Table 1, and Table 2 gives an overview of the different in-silico scenarios. We assume that the porosity equals the extracellular space volume fraction *ϕ* = *ϕ*_PAR_ within the parenchyma and *ϕ* = 1 elsewhere. The effective diffusion coefficient is set equal to that of Gadubutrol in free water in the PVS networks: 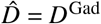;weighted by a dispersion factor *R* in the CSF spaces: 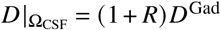; and modulated by the porosity and tortuosity (*λ* = 1.78) in the parenchyma: 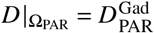. We set *ξ* depending on whether the PVS segment is fully embedded within the CSF spaces (*ξ* = *ξ*_CSF_) or parenchyma (*ξ* = *ξ*_EF_), or neither (Appendix A.1). Additionally, we increase the permeability *ξ* (100×) at the the leaf nodes of Λ_*a*_, linearly decreasing to the baseline permeability at the next network node (≈ 0.4 mm), to account for the transition from explicitly resolved PVSs to their unresolved continuation in the CSF spaces and parenchyma.

**Table 2.** Overview of models and model variations. (− denotes equal to the corresponding baseline value). Each PVS cross-section is modelled as a concentric annulus with inner radius *R*_1_ and outer radius *R*_2_ and such that *R*_2_ = *βR*_1_. Let *ξ* denote the astrocytic endfeet permeability estimated by Koch et al.^91^. The diffusion coefficients *D*^Gad^ represent the (free) diffusion coefficient of Gadubutrol in CSF (water), and 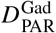 its effective diffusion coefficient in human cortical tissue, all at body temperature^76,109^. *R* is the combined cardiac and respiratory dispersion enhancement field (see Methods). **u**_CSF_ is the CSF velocity field induced by CSF production in Ω_CSF_, and 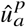 and 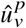 the axial velocity induced in the periarterial and perivenous networks, respectively, by the corresponding pressure differences. 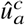 and 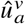 is the periarterial velocity induced by arterial pulse wave wall motion and slow vasomotion, respectively. Last, 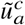 and 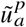 are as 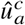 and 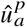, respectively, but with dilated PVS.

| Description | $R_1, R_2$ | $\xi_{\text{PVS-CSF}}$ | $D\vert _{\Omega_{\text{CSF}}}$ | $D\vert _{\Omega_{\text{PAR}}}$ | $\mathbf{u}\vert _{\Omega_{\text{CSF}}}$ | $\hat{u}\vert _{\Lambda_a}$ | $\hat{u}\vert _{\Lambda_v}$ |
| --- | --- | --- | --- | --- | --- | --- | --- |
| baseline | $R_2 = 2R_1$ | $\xi$ | $(1+R)D^{\text{Gad}}$ | $D_{\text{PAR}}^{\text{Gad}}$ | $\mathbf{u}_{\text{CSF}}$ | $\hat{u}_a^p + \hat{u}_a^c$ | $\hat{u}_v^p$ |
| no CSF production | – | – | – | – | $\mathbf{0}$ | $\hat{u}_a^c$ | 0 |
| reduced pulsatility | – | – | $(1+0.1R)D^{\text{Gad}}$ | – | – | – | – |
| increased CSF dispersion | – | – | $(1+10R)D^{\text{Gad}}$ | – | – | – | – |
| higher PVS flow | – | – | – | – | – | $\hat{u}_a^p + \hat{u}_a^c + \hat{u}_a^v$ | – |
| higher PVS exchange | – | $100\xi$ | – | – | – | – | – |
| dilated PVS | $R_2 = 3R_1$ | – | – | – | – | $\tilde{u}_a^p + \tilde{u}_a^c$ | – |

### Numerical approximation, implementation, and verification

We give a concise summary of the numerical approach here (see Supplementary methods for a comprehensive description). To solve (1) numerically, we use the interior penalty discontinuous Galerkin (DG) method with weighted averages and upwinding for the convection term in the 3D domains^113^ – to ensure numerical stability and minimize the artificial presence of negative concentration values. For the 1D domain, we use continuous finite element spaces defined over each of the networks and stabilized with numerical diffusion when 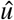 is non–zero. Note that the 3D–1D coupling in (1) leads to a 3D solution *c* with low regularity properties. Thus, one can only expect lower-order error convergence near Λ_*a*_ and Λ_*v*_, see the references^114^ and^105^ for the numerical analysis of DG and continuous Galerkin schemes, respectively. However, away from these 1D networks, almost optimal approximation is expected^115,116^. To numerically solve the Stokes equations to compute the CSF flow velocity and pressure in the ventricular system and SAS, we use finite element spaces that preserve the incompressibility condition on the discrete level^117^. This requirement is important for numerical stability when subsequently solving (1)^118^. The spaces we used for the velocity are continuous along the normal direction; continuity of the tangential component is enforced via interior penalty approaches^117^. Optimal approximation properties are expected for this scheme. These numerical methods were implemented using the FEniCS finite element framework^79^, and the 3D-1D coupling is handled by the extension FEniCS_ii_^80^. The correctness and accuracy of the numerical solutions were verified by a series of numerical verification experiments (Appendix A.5).

### Model validation

As the primary means of model validation, we compare the in-silico predictions of tracer enrichment and clearance against glymphatic MRI studies (Discussion). In addition, we compare auxiliary model quantities (CSF flow rates and pressure differences, dispersion factor estimates, and the shapes and sizes of PVSs) with the current literature (Appendix B.1).

## Acknowledgments

All authors express their gratitude to Prof. Jürgen Reichenbach for generously sharing the multi-modal human MR data, and to Prof. Erlend Hodneland for facilitating this data exchange. M.E.R. acknowledges support from Stiftelsen Kristian Gerhard Jebsen via the K. G. Jebsen Centre for Brain Fluid Research. M.C., R.M., and M.E.R. acknowledge support from the Research Council of Norway (RCN) via grant #324239 (EMIx). Simulations were performed on the Experimental Infrastructure for Exploration of Exascale Computing (eX3), which is financially supported by the RCN under contract 270053. M.K. acknowledges support from the RCN grant #303362. The figures 1A, 1D, 2C, 3C, 5A and 6A were created in Biorender.

## Author contributions statement

M.C., M.K, R.M., and M.E.R. conceived and designed the project. M.C., M.K, R.M., and M.E.R. contributed to software development. M.C., M.K, and R.M. conducted the experiments. M.C. analyzed the results and prepared the figures. M.C. M.K. and R.M and M.E.R wrote the manuscript. All authors edited and reviewed the manuscript.

## A Supplementary methods

These sections provide more detailed descriptions of the mathematical models and numerical approximations considered.

### A.1 Mixed-dimensional transport and flow equations

#### A.1.1 Notation

In terms of geometrical domains, we consider the parenchyma Ω_PAR_ ⊂ ℝ^3^ and CSF spaces Ω_CSF_ ⊂ℝ^3^ with Ω = Ω_PAR_ ∪Ω_CSF_ (Figure 1A, Figure 2A–C). The coordinates in these 3D domains is denoted by *x*. The interface between the parenchyma and CSF spaces is given by ∂Ω_PAR_ ∩ ∂Ω_C_n_SF_ and we separate it as two parts: the surface of the lateral ventricles Γ_LV_ and the remaining pial interface Γ_pia_ (thus ∂Ω_CSF_ into two parts: Γ_SSAS_ represents the lower interface towards the spinal subarachnoid space (SSAS), while Γ_AM_ is the outer ∂Ω_PAR_ = Γ_LV_ ∪ Γ_pia_). The remaining, outer, boundary of Ω_CSF_ is again separated interface towards the arachnoid and dura membranes. Γ_AM_ is further subdivided into its lower and upper parts: Γ_AM−L_ and Γ_AM−U_. The boundary towards the spinal cord is denoted by Γ_SC_ and given by Γ_SC_ = ∂Ω_PAR_ \∂Ω_CSF_.

In addition, we consider two sets of perivascular networks: a periarterial network Λ_*a*_ represented by the (connected) centerlines 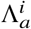 of the arterial tree, and a perivenous network Λ_*v*_ associated with the centerlines 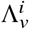 of the veins. Recall that we represent the vascular domains as the union of cylindrical vessels of radius 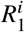 surrounding the centerlines Λ^*i*^. Moreover, we consider the surrounding perivascular spaces as the union of annular cylinders Ω^*i*^ of inner radius 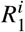 and outer radius 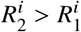 and thus of width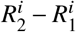, and remark that we interpret 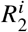 and 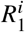 as temporal averages (fixed in time), as the pulsations considered in Appendix A.4 are beyond the temporal resolution of our model. We denote the outer lateral surface of the periarterial and perivenous spaces by Γ_*a*_ and Γ_*v*_, respectively. We omit the subscript *a* or *v* when referring to any such network, vessel segment or perivascular outer surface. We assume that Λ is parametrized by the coordinate *s*, and with a minor abuse of notation, simply write *s ∈* Λ to represent the point *λ*(*s*) on Λ corresponding to *s*. Finally, we view each perivascular network both as a geometric domain and as a directed graph with the centerlines {Λ^*i*^} as oriented edges and the connections as nodes 𝒱.

#### A.1.2 Solute transport and exchange in intracranial domains (3D) and perivascular networks (1D)

In this section, we describe the mathematical model (1), the associated definitions, interface and boundary conditions in further detail. We refer to the main text for explicit parameter values while giving the general, abstract form here, and to the reference^65,105^ for the derivation and analysis of this 3D-1D model. Recall that we model a concentration field *c* = *c*(*x, t*) for *x ∈* Ω and *t* > 0 defined in Ω_PAR_ and Ω_CSF_ separately. In each of these domains, *c* satisfies

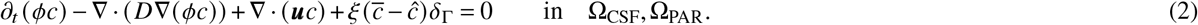

In (2), *ϕ* is the fluid volume fraction (also known as the porosity) defined in the parenchyma (*ϕ* ≪1 in Ω_PAR_) and in the CSF spaces (*ϕ* = 1 in Ω_CSF_). In the parenchyma, *ϕ* represents the extracellular space volume fraction, and thus *c* here generally represents the intrinsic (in contrast to the superficial) concentration^76^. Moreover, *D* is the effective diffusion coefficient of the relevant solute in the respective media which takes different values over the CSF spaces and the parenchyma, depending on tortuosity^76^ and dispersive effects; and *u* is a convective velocity field representing the flow of CSF in Ω_CSF_ and the flow of ISF in Ω_PAR_. *ξ* models a transfer or exchange parameter between the 3D domain (Ω_PAR_ or Ω_CSF_) and the perivascular networks Λ_*a*_, Λ_*v*_. To summarize,

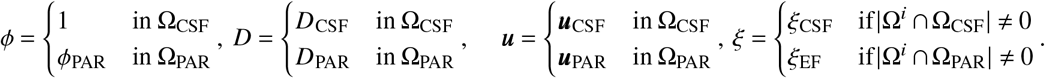

In the above, *ξ* is defined segment-wise: for each centerline Λ^*i*^ with surrounding PVS Ω^*i*^, |Ω^*i*^ ∩ Ω_PAR_ | (resp. |Ω^*i*^ ∩ Ω_CSF_ |) is nonzero whenever Ω^*i*^ intersects Ω_PAR_ (resp. Ω_CSF_). If the surrounding PVS intersects both, then we set *ξ* = *ξ*_EF_ if Ω^*i*^ mainly (80 percent) intersects Ω_PAR_, in which case the interaction with Ω_CSF_ is ignored; otherwise, we set *ξ* = *ξ*_CSF_.

Also in (2), the notation 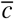 denotes a lateral average of the concentration over the outer perivascular surfaces, defined for each centerline Λ^*i*^ and each point *s ∈* Λ^*i*^ by

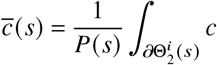

where 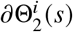 *s* is the outer boundary of the cross-section Θ *s* of the PVS Ω^*i*^ at *s* and *P s* is the perimeter of Θ *s*. Moreover, for both the periarterial and perivenous networks (Λ_*a*_, Λ_*v*_ with outer PVS boundaries Γ_*a*_, Γ_*v*_), the term *δ*_Γ_ is concentrated on the outer lateral surfaces of the PVSs, and defined in terms of its action on any sufficiently smooth function *v* : Ω → ℝ such that

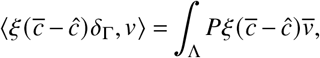

under the assumption that *ξ* is constant on each 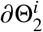.

On the interface between the brain and the CSF spaces Γ_pia_ ∪ Γ_LV_, we prescribe the following interface conditions, which represent a semi–permeable interface, writing *c*|_ΩPAR_ = *c*_PAR_ and *c*|_ΩCSF_ = *c*_CSF_:

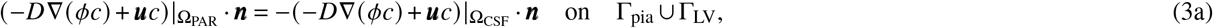

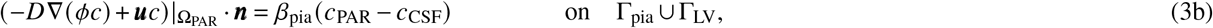

where *n* = *n*_PAR_ is the normal vector field defined over the interface, outward-pointing when viewed from Ω_PAR_, and *β*_pia_ ⩾ 0 is a permeability constant.

We supplement (2) and (3) with the following boundary conditions representing a given molecular influx at Γ_SSAS_, molecular efflux at a constant update rate *β*_exit_ at Γ_AM−U_, and no influx or efflux elsewhere from the CSF spaces.

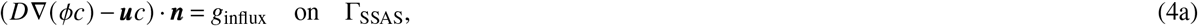

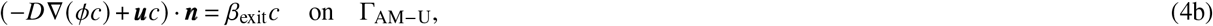

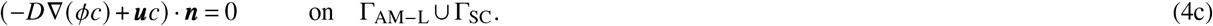

where again *n* denotes the outward-pointing boundary normal.

The concentration 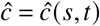 in the periarterial and perivenous networks (*s ∈*Λ_*a*_, *s ∈*Λ_*v*_), entering in (2), represents the concentration in the perivascular space averaged over each perivascular cross-section and is governed by

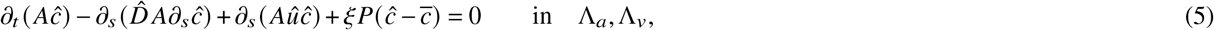

where in Λ_*a*_, Λ_*v*_ refers to in each Λ^*i*^ in each of these networks. In (5), *A* = *A ∈ s* is defined as the PVS cross-sectional area; i.e., for each *s* ∈ Λ^*i*^ with associated PVS Ω^*i*^, the area of the cross-section Θ^*i*^ *s*. Also, 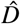 is the effective diffusion coefficient, and 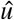 is a convective velocity representing an (average) CSF flow velocity in the axial direction of the perivascular spaces.

To complete (5), we prescribe bifurcation conditions at internal nodes of the perivascular networks Λ_*a*_, Λ_*v*_ and boundary conditions at the end nodes. To this end, define the set of internal nodes ℬ ⊂𝒱as the set of nodes that are connected to two or more edges and the set of end nodes as 𝒩 ⊂𝒱, write 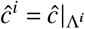, and for *y* let *y∈*ℬ let ℰ (*y*)denote the set of (two or three) edges in {Λ^*i*^} sharing the node *y*. To impose continuity of the concentrations, we set that the concentrations when viewed from each edge must match at nodes:

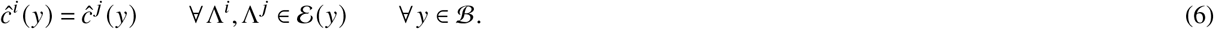

Moreover, to ensure mass conservation, we set that the flux going in and out at nodes should add to zero, or more precisely, that for each *y ∈* ℬ:

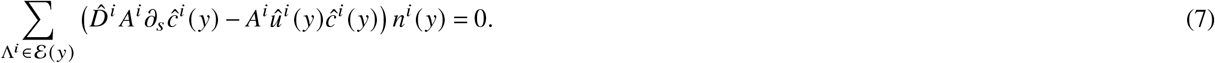

Here the (normal “vector”) function *n*^*i*^ takes the values in {−1, 1} and defines an orientation of vertices of edge Λ^*i*^. Specifically, for an edge Λ^*i*^ oriented from *y*_in_ to *y*_out_ we set

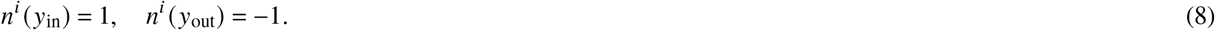

Finally, we impose a homogeneous Neumann condition at all network end nodes to augment (5):

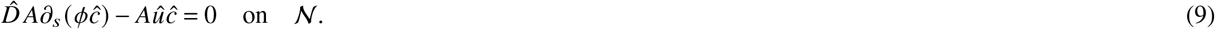

This no-flux condition states that the network end nodes represent barriers for molecular efflux into or out of the perivascular network, and thus that all solute exchange between the PVSs and their surroundings takes place via the lateral outer PVS surface and is regulated by the exchange parameter *ξ* cf. (2) and (5). Larger particles have been observed to accumulate within the PVS as the surface arteries penetrate into the brain parenchyma^32,33^, and (9) is appropriate to represent such behavior. However, to represent a continuously extending PVSs also along penetrating vessels, this condition should be revisited.

#### A.1.3 Modelling CSF flow via the incompressible Stokes equations

The time-independent, incompressible Stokes equations model the flow of an incompressible Newtonian fluid at low Reynolds numbers, and read as follows: over the domain Ω_CSF_ ⊂ ℝ^3^, find the velocity vector field *u* : Ω_CSF_ → ℝ^3^ and the pressure field *p* : Ω_CSF_ → ℝ such that

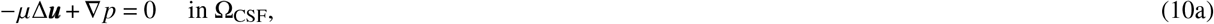

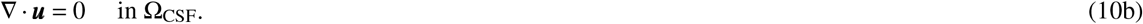

In addition, we impose the following boundary conditions to model flow induced by CSF production in the choroid plexus with Γ_AM−U_ as the main efflux site:

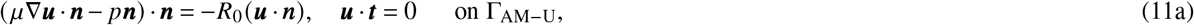

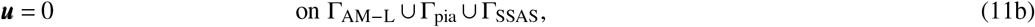

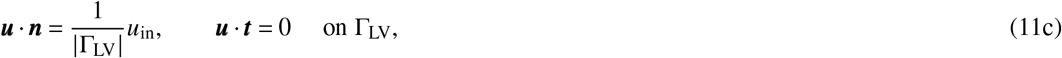

where *n* and *t* denote the unit outward normal and tangent vectors to the boundary respectively, *µ* is the (dynamic) CSF viscosity, *R*_0_ > 0 is a resistance parameter for CSF efflux, and *u*_in_ is a given fluid influx, here across the lateral ventricle wall Γ_LV_. We consider alternative variations of these boundary conditions in connection with estimating dispersion effects induced by CSF pulsatility, see Appendix A.3.

#### A.1.4 Steady flow in perivascular networks induced by pressure differences

To model steady flow induced by pressure differences between end nodes in a perivascular network Λ, with edges {Λ^*i*^} _*i*_, internal nodes ℬ and end nodes 𝒩, we consider the following system of hydraulic network equations^119–121^. The unknowns are the PVS flux 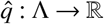 and pressure 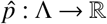, which represent the fluid flux and the average pressure across cross-sections of the PVSs, respectively. These are defined for each Λ^*i*^ by solving:

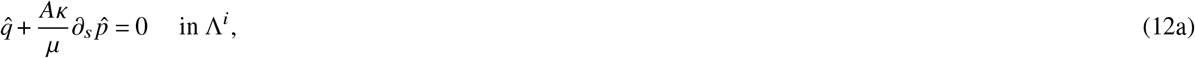

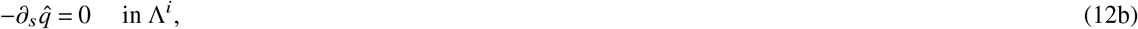

where *A* is the area of the PVS cross-sections, *µ* is the CSF viscosity as before, and κ is derived from an assumption of Poiseuille flow in the annular cross-section of the PVS as^119^:

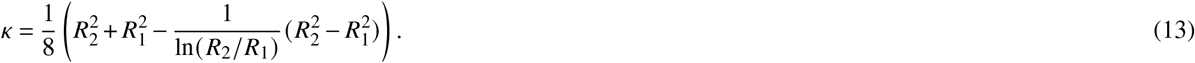

The PVS flux 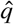 and the associated average PVS velocity in the axial direction 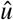 are directly related by the cross-sectional area

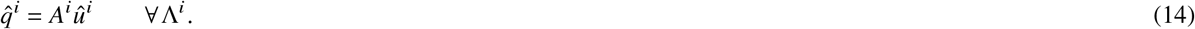

In addition, to complete (12), we impose continuity of the fluid pressure and conservation of flux at each internal node. Write 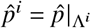. These conditions then read as: for each *y ∈* B with connected edges ℰ (*y*):

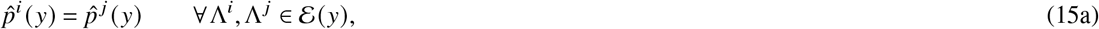

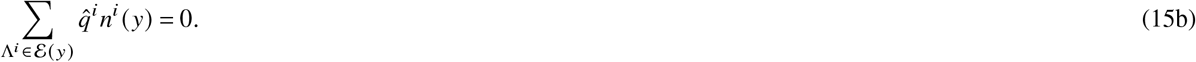

where *n*^*i*^ is as defined by (8). Finally, to drive this flow, we impose given fluid pressures *p*_0_ at the network end nodes:

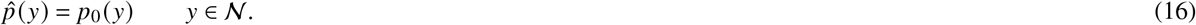

In the case where modelling flow in the perivascular networks Λ_*a*_, Λ_*v*_ induced by CSF production, we impose the fluid pressure *p*_CSF_ computed in the CSF spaces Ω_CSF_ (and its harmonic extensions into Ω_PAR_) as the given fluid pressures *p*_0_. Note that we here consider steady PVS flow in a non-moving domain; the effect of domain motion is addressed in Appendix A.4.

### A.2 Numerical solution of the transport and flow equations

We solve the models presented in Appendix A.1 numerically using finite element methods. The following subsections provide more details, model-by-model.

#### A.2.1 Computational mesh entities: definitions and common notation

We consider a mesh 𝒯= {*E*} of Ω = Ω_PAR_ ∪ Ω_CSF_, consisting of tetrahedral mesh cells *E*, and conforming to the domains Ω_PAR_ and Ω_CSF_ and to the CSF-brain interface Γ_pia_ ∪ Γ_LV_. We denote the restriction of 𝒯to Ω_PAR_ and Ω_CSF_ by 𝒯_PAR_ and 𝒯_CSF_, respectively. The collection of all interior facets (i.e. triangular faces of the tetrahedral mesh cells) in 𝒯_PAR_ and 𝒯_CSF_ are denoted by ℱ_*i*,PAR_ and ℱ_*i*,CSF_, respectively. We define the union of facets interior both to 𝒯_PAR_ and 𝒯_CSF_ as ℱ_*i*_ = ℱ_*i*,PAR_ ∪ ℱ_*i*,CSF_.

#### A.2.2 Finite element solution of coupled 3D-1D solute transport equations

We consider the system of coupled 3D-1D solute transport equations given by (2) and (5) with the interface conditions (3), the bifurcation conditions (6) and (7), and the boundary conditions (4) and (9). We discretize these equations using an implicit finite difference scheme in time, a discontinuous Galerkin (DG) finite element method with upwinding in space for the 3D domain to accurately capture sharp boundary layers, and a continuous Galerkin method for the 1D networks. We remark that the transport in the CSF spaces is highly convection dominated, with an average Péclet number of 402 and maximum of 9542 (assuming a characteristic length of 10 cm, and accounting for the increased diffusivity due to dispersion).

We first consider the discretization of (2) and introduce the discrete space

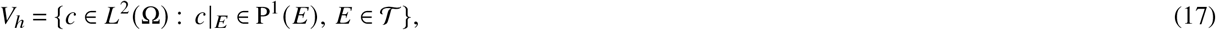

where *L*^2^ (Ω) is the space of square-integrable functions defined over Ω and P^1^ (*E)* denotes the space of polynomials of total degree ⩽ 1 defined over the tetrahedra *E*. To discretize the diffusion term in (2), we use a symmetric weighted interior penalty DG formulation, referring to^113^ and [122, Section 4.5.2.3] for details on this method. Recall that *ϕ* is constant in each domain Ω_PAR_ and Ω_CSF_, and thus in particular that ∇(*ϕc*) = *ϕ*∇*c* on each *E ∈* 𝒯. Define for 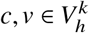:

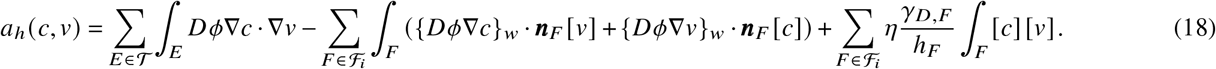

In (18), for each facet *F ∈* ℱ^*i*^ shared between cells *E*^1^ and *E*^2^, we associate a facet normal vector *n*_*F*_ pointing from *E*^1^ to *E*^2^. The facet diameter is denoted by *h*_*F*_, the jump [·] is given by 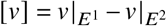, and the unweighted average {·} and weighed average {·}_*w*_ are defined as:

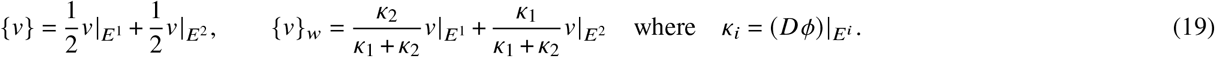

The parameter *γ*_*D,F*_ is the harmonic mean of the porosity-weighted diffusion coefficient given by

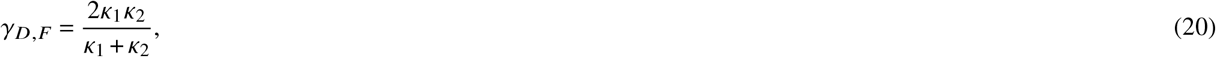

and *η* is a user-specified penalty parameter (we set *η* = 1000). To discretize the convection term in (2), we use upwinding, see [122, Section 2.3.1] and the references therein. For 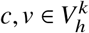, define

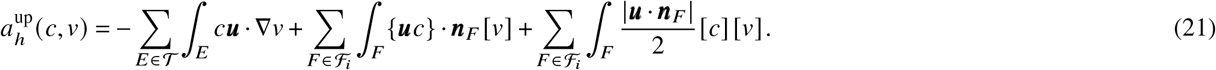

Our discrete formulation for (2) with the interface conditions (3) and boundary conditions (4) and given initial conditions 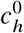 then reads: for *n* = 1, 2, …, with *t*^*n*^ − *t*^*n*−1^ = *τ*, find 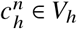 such that for all *v ∈ V*_*h*_:

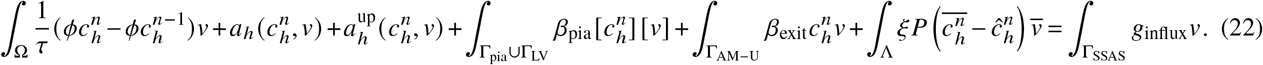

To discretize (5) with the bifurcation conditions (6) and (7), and the boundary conditions (9), we use the space of continuous piecewise linear polynomials defined over Λ:

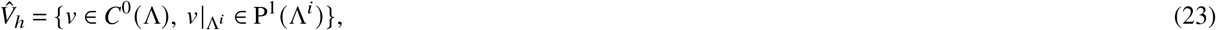

where P^1^ (Λ^*i*^) is the space of linear polynomials on each Λ^*i*^. The discrete formulation then reads: for *n* = 1, 2, …, find 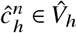 such that for all 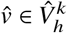:

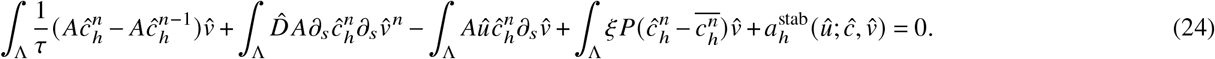

In the above, if 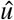 is non–zero on Λ^*i*^, then the artificial diffusion 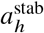 stabilization term is nonzero and is given below, for more details see for [123, Section 12.6].

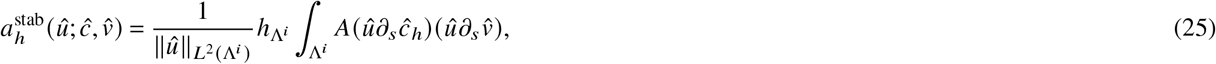

where 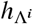 is the mesh–size of Λ^*i*^. Note that the condition (7) is enforced weakly in the above formulation.

*Summary*. The discretization for (2) and (5) complemented with boundary conditions ((4) and (9)), interface and bifurcation conditions ((3), (15), and (6)), and intial conditions 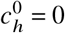 and 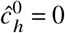 is the following:

For *n* = 1, 2, …, find 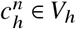 and 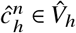 such that the coupled equations (22) and (24) hold for all *v ∈ V*_*h*_ and for all 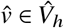.

#### A.2.3 Finite element solution of the incompressible Stokes equations

We consider an H(div)-based finite element approximation of the incompressible Stokes equations (10) defined over Ω_CSF_ with the boundary conditions (11). Following^117^, we approximate the velocity field *u* and the pressure field *p* with the following finite element spaces:

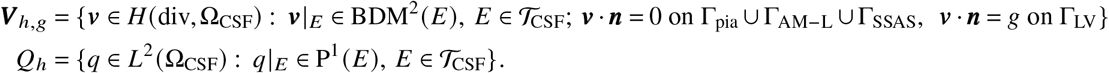

Here, *H* (div, Ω_CSF_) is the space of *L*^2^ (Ω_CSF_) vector fields with *L*^2^ (Ω_CSF_) divergence, BDM^2^ is the Brezzi-Douglas–Marini element^124^ of degree 2, *g* is a given constant, and *n* is the unit outward normal vector to each facet. Given any vector *v*, the normal and tangential components on each facet are denoted and given by

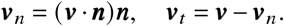

Since *V* _*h,g*_ ⊂ *H*(div, Ω_CSF_), then [*v*_*n*_] = 0 on ℱ_*i*,CSF_, the interior facets to Ω_CSF_. Continuity in the tangential component is enforced weakly via interior penalization. For convenience, we collect all facets exterior to the CSF space Γ_AM−U_ in the set

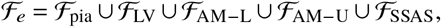

where facets lying on the pial interface Γ_pia_ are denoted by ℱ_pia_, on the lower and upper outer (arachnoid) boundary Γ_AM−L_ and Γ_AM−U_ by ℱ_AM−L_ and ℱ_AM−U_, respectively, on the boundary towards the spinal SAS Γ_SSAS_ by ℱ_SSAS_, and on the surface of the lateral ventricles by ℱ_LV_. Now, define the form

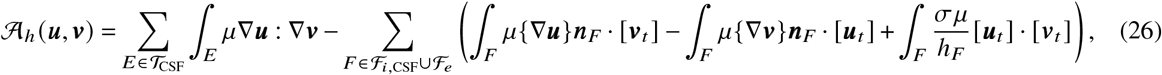

where on exterior facets the average and jump operators take the one-sided values. We set the penalty parameter for the tangential continuity to be *σ* = 20. The finite element discretization of the incompressible Stokes equations is then to find (*u*_*h*_, *p*_*h*_) *∈ V* _*h,g*_ ×*Q*_*h*_ with 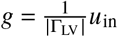 such that

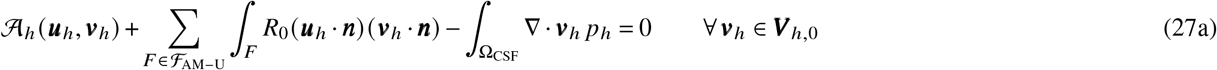

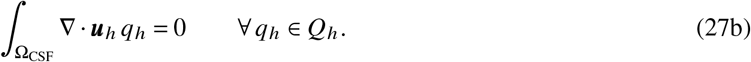

#### A.2.4 Finite element solution of the perivascular network equations

We consider meshes ℐ_*a*_, ℐ_*v*_ representing a conforming subdivision of each of the perivascular networks Λ_*a*_, Λ_*v*_. Relative to each mesh ℐ, we define the space of (discontinuous) piecewise constants 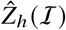 and define spaces of continuous piecewise linears 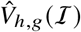 with prescribed boundary node values (on 𝒩) given by *g*:

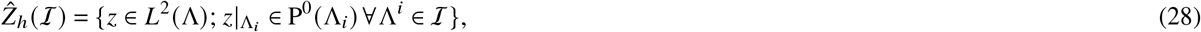

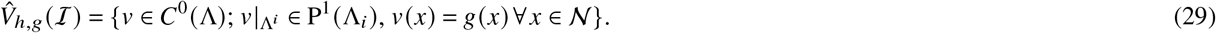

To discretize (12) with the bifurcation conditions (15) and boundary conditions (16), we use the space of continuous functions 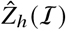 to enforce the continuity of the pressure at bifurcation points, while the conservation of flux is enforced (weakly) through the variational formulation. The discrete variational form of the equations then reads: Find 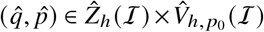 such that

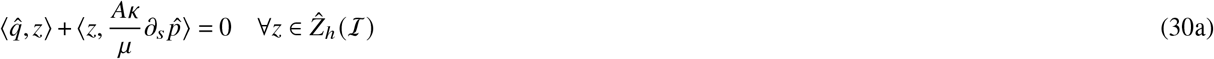

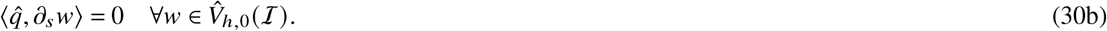

where ⟨·, ·⟩ denotes the *L*^2^(ℐ) -inner product and defined segment-wise. For a stability and convergence analysis of this discrete model, we refer to the reference^121^.

### A.3 Estimating dispersion factors from pulsatile CSF flow

The cardiac (*∼*1Hz) and respiratory (*∼*0.25Hz) cycles induce pulsatile flow of CSF in the ventricular system and in the cranial and spinal SAS. Pulsatile flow leads to dispersion which in turn may enhance molecular transport^42,43,46,77,106,107^. To account for the dispersive effects over a longer time scale (hours to days), and in the absence of measurements or estimates of dispersion coefficients in human CSF spaces, we adapt existing theoretical estimates^43,107^. More specifically, we compute spatially-varying dispersion enhancement fields *R*_*c*_ and *R*_*r*_ (Figure 2G, I), associated with the cardiac and respiratory cycles respectively, via the algorithm presented below. These fields then contribute to the diffusion coefficient in Ω_CSF_ in (2) as *D* = (1 + *R*_*c*_ + *R*_*r*_) *D*^Gad^.

i. To account for viscous forces, we compute spatially-varying CSF pressure fields 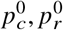 in Ω_CSF_ corresponding to the Stokes flow induced by the peak volumetric reduction of the CSF space in the respective cycle (Figure 2F, H). More precisely, we numerically solve the incompressible Stokes equations (10) equipped with the following boundary conditions mimicking a dilation of the brain parenchyma with the spinal SAS as the only route for CSF efflux:

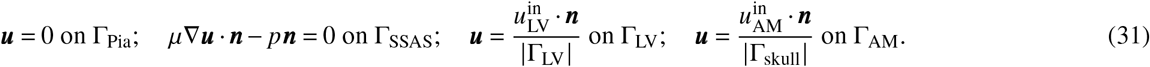 In the cardiac cycle case, we set 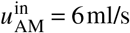^56,81^ and 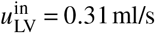^55^ to solve for 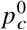 In the respiratory cycle case, we set 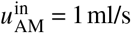^82^ and 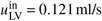^83^ to solve for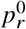.
ii. We also estimate the Womersley numbers *α*_*c*_, *α*_*r*_ associated with the cardiac and respiratory flow patterns, respectively, by the definition

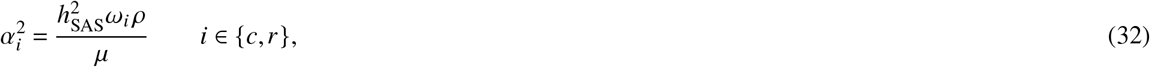

with CSF density *ρ* = 10^3^ kg/m^3^, a mean CSF space width *h*_SAS_ = 1.5mm, and CSF viscosity *µ* given in Table 1. For the cardiac cycle, we consider an angular frequency *ω*_*c*_ = 2*π*, while for the respiratory cycle, we set *ω*_*r*_ = 0.5*π*. The resulting (square) Womersley numbers are 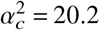 for the cardiac cycle and 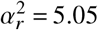 for the respiratory cycle.
iii. To account for inertial forces in addition to the viscous forces, we use the Womersley numbers *α*_*c*_, *α*_*r*_ to calculate upscaled pressure fields 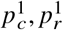 from 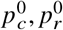 as:

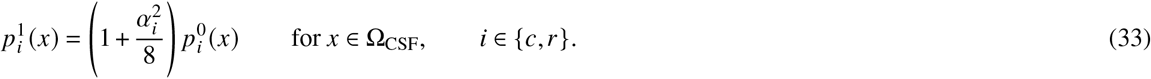 Note that this scaling is based on theoretical considerations on the ratio of oscillatory flow to steady flow impedances in a tube [125, Chap. 4.3.].
iv. Further, assuming unsteady dispersion, we follow Sharp et al.^43^ to estimate local enhancement factors *S*_*c*_, *S*_*r*_ from the non-dimensionalized pressure gradients:

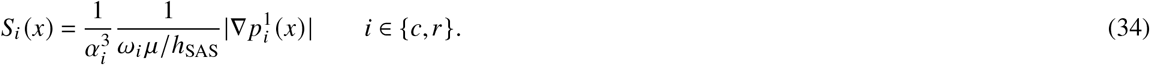
v. Finally, we define the cardiac and respiratory dispersion enhancement factors *R*_*c*_ and *R*_*r*_ by smoothing *S*_*c*_ and *S*_*r*_, respectively, to account for the non-local nature of dispersion. Specifically, for *i∈* {*c, r*}, we define *R*_*i*_ : Ω_CSF_ →ℝ^+^ by solving a heuristic weighted Helmholtz problem over Ω_CSF_ with *S*_*i*_ as the right-hand side:

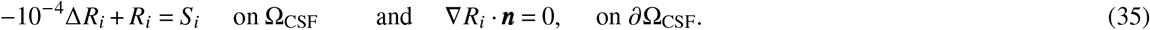

Considering the uncertainty associated with the validity of simplifying assumptions, the resulting estimates of the cardiac and respiratory dispersion factors *R*_*c*_ and *R*_*r*_ should be viewed as heuristic rather than absolute.

### A.4 Estimating net perivascular flow induced by peristaltic waves

We use the theoretical framework previously introduced by Gjerde et al.^40^ to compute an analytic estimate of the time-averaged (or *net*) flow rates 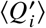 (mm^3^/s) induced by peristaltic pumping in a perivascular network Λ =∪ _*i*_Λ_*i*_. The motion of the (inner) vascular wall is described by a periodic traveling (peristaltic) wave of relative amplitude *ε*, wave length *λ* (mm) and frequency *f*, acting normal to the wall. By definition, the wave number is *k* = 2*π*/*λ* and the angular frequency is *ω* = 2*π f*. Each PVS segment Λ_*i*_ has length *L*_*i*_ with wave-relative length *ℓ*_*i*_ = *k L*_*i*_, baseline inner radius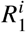, fixed outer radius 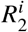, and outer-to-inner ratio 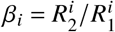. These geometric parameters and the assumption of annular cylindrical PVS segments yield hydraulic resistances ℛ_*o,i*_ and additional characteristic parameters, see^40^ for the complete definitions and schematics. Since the analytical estimate is derived under the assumption that *k L*_*i*_ ≈ 𝒪(1)^40^ and has been verified against numerical simulations for *k L*_*i*_ of the order 10^−1^ − 10^2^ [40, Table I], we consider it applicable for the wave lengths and vascular network data considered here in which *k L*_*i*_ range from 0.15 to 30 for the strong vasomotion and 0.0015 to 0.30 for the cardiac waves (Table 3).

**Table 3.**
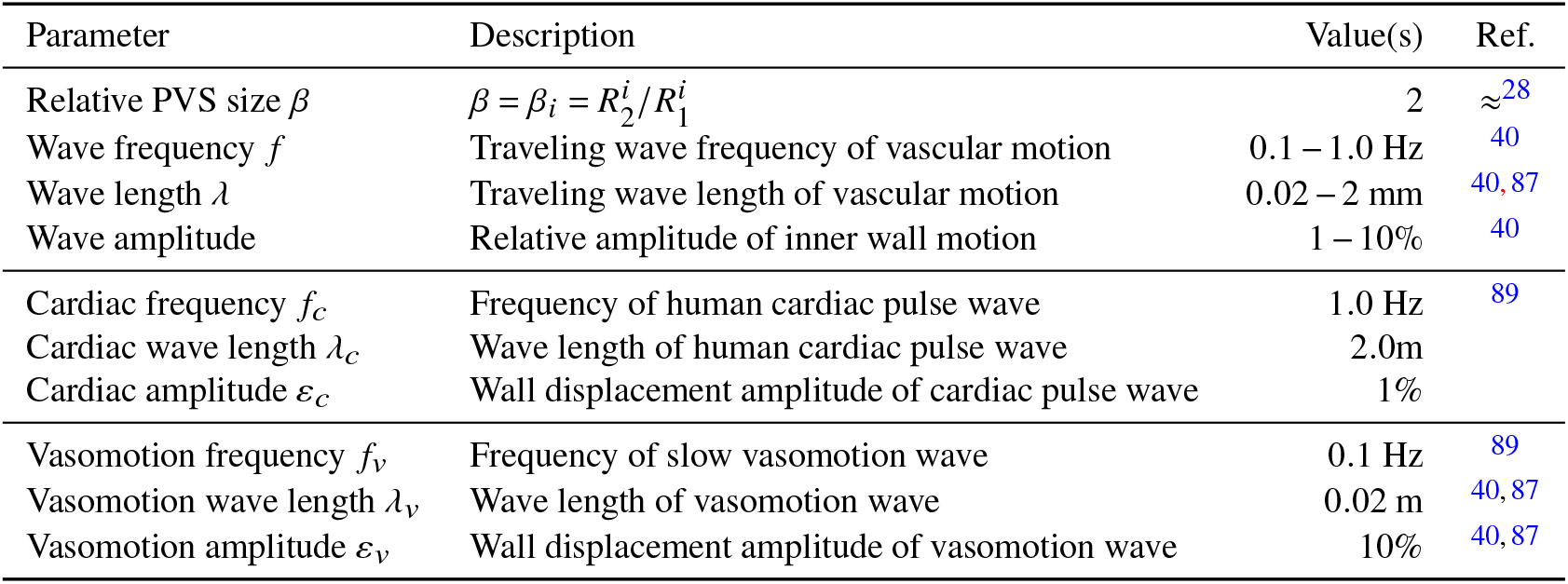
Perivascular flow induced by vascular wall motion: overview of parameters.

This theoretical formalism^40^ is defined relative to a network in the form of a directed, bifurcating tree with a single supply node/root *i*_0_. To extend to a network of cerebral arteries with multiple supply nodes (such as the basilar and two internal carotid arteries in the current data set^66^), we separate the network Λ into edge-disjoint subnetworks Λ^1^, Λ^2^, Λ^3^, one for each of the supply nodes (Figure 1A, C, Figure 7). Each node is assigned to the subnetwork associated with the nearest supply node, and edges between nodes are preserved (Figure 7B). Next, we compute a minimal, bifurcating and directed tree representation of each subnetwork: 𝒯^1^, 𝒯^2^, 𝒯^3^ (Figure 7C). Each tree 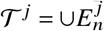 consists of the subset of the nodes from Λ^*j*^ that have degree 1 (are leaf or root nodes) or degree 3 (are true bifurcation points), and each path between nodes with degree ≠ 2 in Λ^*j*^ is represented in 𝒯^*j*^ by an edge *E*^*j*^ with edge length *L* corresponding to the total length of the original path and edge radius *R*^*j*^ as the average of the path radii.

**Figure 7.**
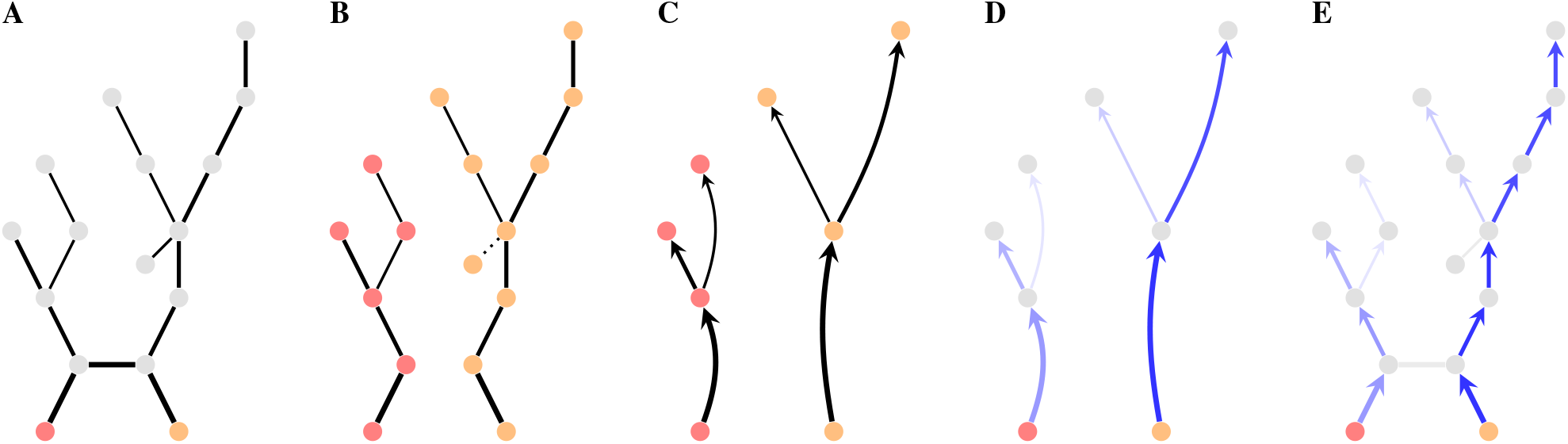
Estimating the perivascular flow induced by vascular wall motion. For a vascular network with vessels represented by edges of varying radii and length connected at nodes (**A**), and one or more supply nodes (here two, marked in red and orange), we compute one subnetwork for each supply node by proximity (**B**). Each subnetwork is reduced to a minimal, bifurcating and directed tree while preserving path lengths and averaging radii (**C**) which is then used to compute the net flow induced by the peristaltic wave in each branch of the minimal subnetworks (**D**). Finally, we distribute the computed flow onto the original segments (**E**).

For each subtree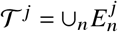, we compute the time-averaged downstream flow rate 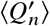 induced by the vascular wall motion for each edge *n* via [40, eq. (5), (34)] (Figure 7D). We next assign this flow rate 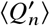 to each of the branches Λ_*i*_ that form the path 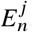, thus yielding 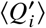 for each perivascular segment Λ_*i*_ while ensuring that mass is conserved (Figure 7E).

For the segment(s) ignored in the separation step (Figure 7B), we set a flow rate of zero. Finally, we define the mean longitudinal perivascular velocity induced by the peristaltic wave 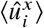 by dividing the flow rate 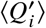 by the cross-section area

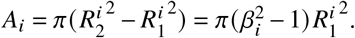

### A.5 Numerical verification

We assess the numerical accuracy and convergence of our simulation results by performing a series of experiments with different spatial and temporal resolutions. Specifically, we generate a sequence of three meshes (Figure 8) with an increasing number of computational mesh vertices and cells, solve all relevant simulation steps (CSF flow and intracranial transport computations) for the baseline model on each mesh, and compare the results across meshes with respect to a set of key quantities of interest. The mesh refinement employed here is localized near expected sharp concentration gradients. This allows us to better capture dynamics and reduce undershoots, see Figure 9. Similarly, we investigate the effect of the time step size by solving the intracranial transport model on the standard resolution mesh with different time steps: 1, 2, and 4 minutes.

**Figure 8.**
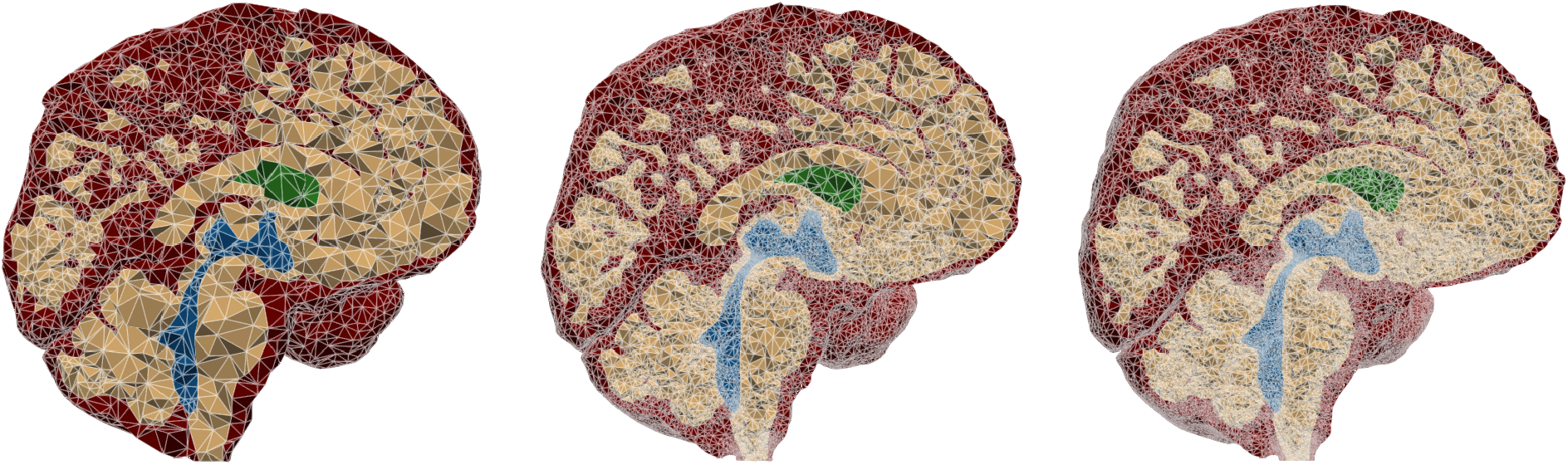
Illustration of the three different meshes; from left to right: low resolution, standard resolution, high resolution.

**Figure 9.**
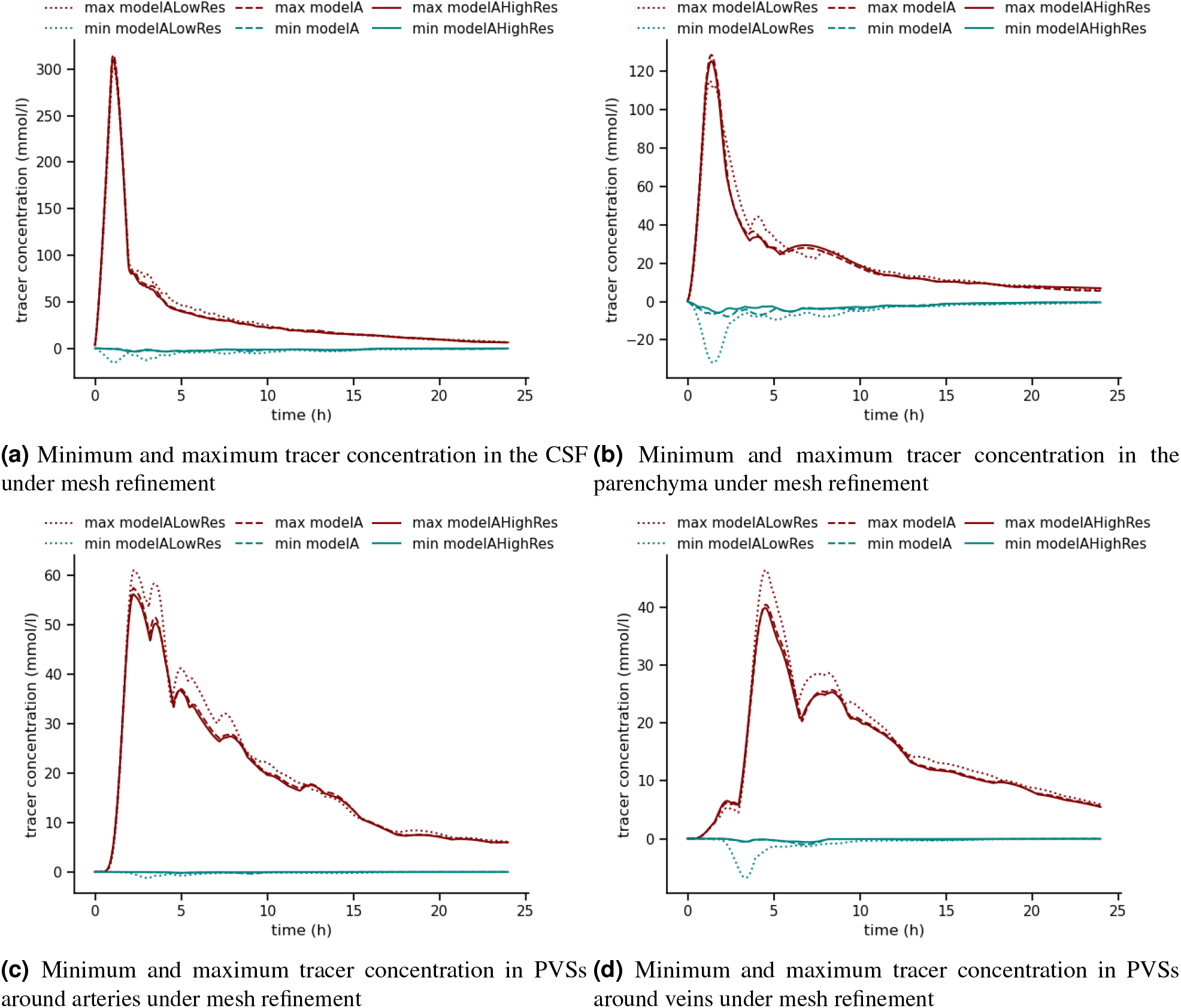
Minimum and maximum tracer concentrations over the first 24 h after injection on the CSF and parenchyma (a), and the arterial and venous PVS (b) computed on the low resolution (LowRes), standard and high resolution (HighRes) meshes with a timestep of 2 min for the baseline model (Model A).

Considering the mean tracer concentrations in the CSF, parenchyma and arterial and venous PVS domain over the first 24 h after injection, we observe negligible changes with both mesh and time refinement (Figure 9 and Figure 11). As an additional verification step, we compute the mean and maximum dispersion enhancement factor, the maximum CSF pressure and velocity in both the cardiac-driven and CSF production-induced flow fields, and mean concentrations at 3, 6, 12 and 24 h for all mesh resolutions and time steps (Figure 10). While the maximum dispersion factor increases by about 60 % from the low resolution to the standard mesh, it stabilizes with the next refinement step. All other quantities change with less than 10 % with mesh refinement, and less than 1 % with time refinement. We thus conclude that the standard resolution mesh and a time step of 2 min offer sufficient accuracy for our simulations, and remark that all reported results are obtained with the standard resolution mesh.

**Figure 10.**
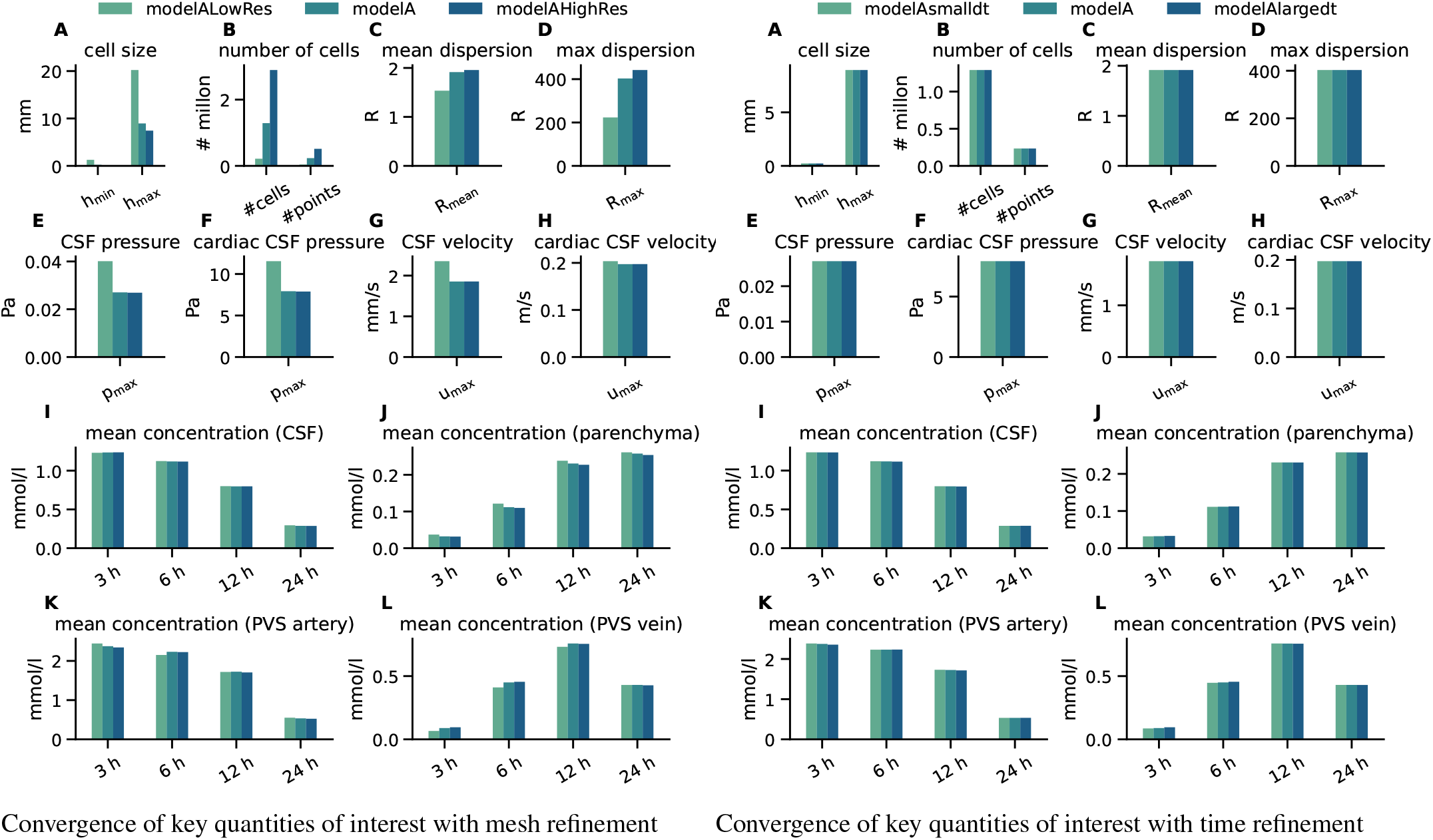
For both left and right panels: A: Minimal (h_min_) and maximal (h_max_) mesh cell sizes (computed as cell circumradius × 2); B: number of mesh vertices and tetrahedral cells in each mesh; C: mean cardiac dispersion enhancement factor *R*; D: maximum cardiac dispersion enhancement factor *R*; E: maximum pressure in steady CSF production flow; F: maximum pressure in cardiac-driven CSF flow; G: maximum CSF velocity in steady CSF production flow; H: maximum CSF velocity in cardiac-driven CSF flow; I–L: mean tracer concentration in the CSF, parenchyma, arterial PVS and venous PVS after 3, 6, 12 and 24 hours.

**Figure 11.**
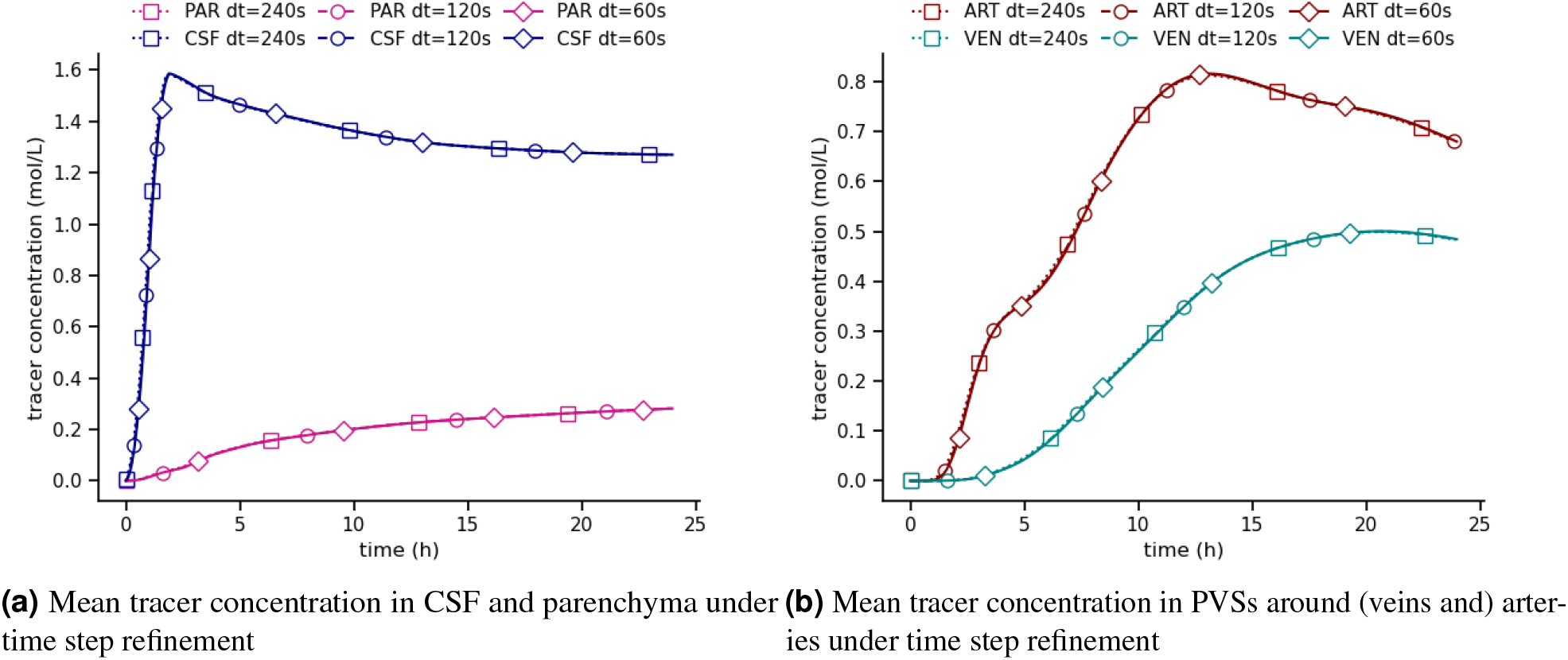
Mean tracer concentrations after up to 24 h in the CSF and parenchyma (a), and the arterial and venous PVS (b) computed on the standard resolution mesh for timesteps of 1, 2, and 4 minutes (dt of 60, 120, or 240 seconds)

Finally, to check for numerical mass conservation, we perform an additional simulation not allowing for tracer efflux across the outer boundary, and confirm that the total amount of tracer is preserved over time after the initial influx phase (Figure 12).

**Figure 12.**
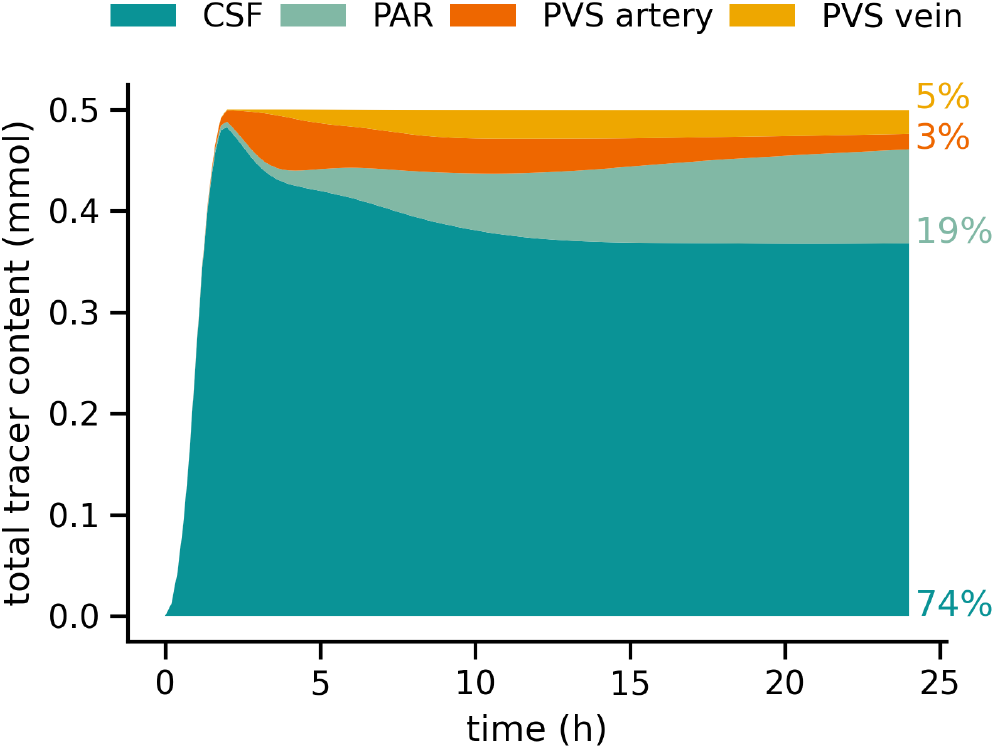
Total tracer content in the CSF, parenchyma, and arterial and venous PVS for a variant of the baseline model without tracer outflow. The total amount of tracer is constant after the initial influx phase demonstrating that the numerical scheme conserves mass globally.

## B Supplementary discussion

### B.1 Extended model validation

In addition to the comparison of our in-silico predictions of tracer enrichment and clearance against glymphatic MRI, we here compare auxiliary model quantities against the literature as additional model validation.

#### CSF flow and pressures in the SAS and ventricular system

The dynamics of human CSF flow and pressure are better quantified, by way of clinical imaging, in-vitro studies, and computational modelling, in other areas of the ventricular system^53,55,56,78,126–128^. Linninger et al^126^ model CSF flow and pressure dynamics induced by CSF production and cardiac pulsatility under normal and hydrocephalic conditions, and report of very good agreement with Cine (phase-contrast) MRI measurements. Our estimates of the maximum intracranial pressure difference, 10 Pa from the cardiac contribution and 26 mPa from CSF production, is in perfect agreement with their maximum transmantle pressure difference of *∼* 10 Pa, and also in very good agreement with mean pressure differences of 11.5 Pa measured clinically between sensors placed subdurally and in the lateral ventricle^55^. Liu et al^128^ report of cardiac and respiratory pressure differences across the aqueduct of 12.1 ± 5.7 Pa and 9.5± 7.2 Pa, respectively; thus our baseline estimate of the respiratory contribution 1.4 Pa may be an underestimation. On the other hand, our cardiac- and respiratory-driven CSF flow estimates peak at 19.8 cm/s and 4.8 cm/s in the caudal direction, respectively, which are higher than phase-contrast MRI measurements of cardiac and respiratory CSF flow components^129,130^. Some variation in CSF flow velocities is expected considering that the values from MRI represent averages^130^ and that the geometry of the CSF spaces strongly affects peak velocities^55,56^. Hornkjøl et al^78^ model the flow dynamics induced by CSF production in the choroid plexus and report a peak CSF velocity of 8.9 mm/s in the aqueduct, which is 4.8 × higher than our values of 1.85 mm/s. Given that we use the same production rate, this deviation again illustrates the impact of potential differences in the (aqueduct) geometry on local velocities.

#### Dispersion in the SAS, ventricular system and PVS

This pulsatile flow of CSF in the SAS, ventricular system and PVSs leads to an increase in effective solute diffusivity^42,43,77,131,132^ via a process known as Taylor dispersion^106,107^. Previous estimates of the magnitude of this effect in the CSF spaces vary significantly: from an enhancement factor of 0.05–1 in periarterial spaces surrounding penetrating arteries^42,46^, to 5–100 in the spinal subarachnoid space^43,131,132^, and up to more than 10000 in surface periarterial spaces^43,77^. The large variability can (at least partly) be attributed to methodological differences; e.g. different assumptions on the medium, domain width, pressure differences and/or fluid velocities, the diversity of CSF flow characteristics, as well as a high likelihood of spatial variations. Hornkjøl et al^78^ consider model variations with constant dispersion factors from 1 up to 1000, and indicate that a value of 10 gives the better agreement with the clinically observed enrichment. Our spatially-varying estimates of the dispersion enhancement factors *R*_*c*_, *R*_*r*_ (with *D* = (1+ *R*_*c*_ +*R*_*r*_) *D*^Gad^) range from 0 to 200 for the cardiac contribution *R*_*c*_ and 0 to 320 for the respiratory contribution *R*_*r*_; and is thus compatible within the previously reported spectrum.

#### Shapes, sizes and structures of the PVS

The shapes, sizes and structures of the PVSs likely vary between species (e.g. mice vs. humans), between spatial compartments (e.g. surface vs. parenchymal), between vessel types (arteries vs. arterioles vs. veins), and in pathologies^23,25,28,32,33,62,64,133–135^. In terms of shape, the PVSs are commonly represented as annular (elliptic) cylinders, though it is well recognized that this represents an idealization^33,64,134–137^. In terms of sizes, Raicevic et al^134^ note that the variation in PVS area is larger between PVS segments than along a single PVS segment and that the PVS area increases with lumen area. In mice, reports of the ratio between PVS and lumen area range from ≈0.35–0.43^64^ up to ≈1.12–1.4^33,134^. In humans, the PVS may be as wide as the associated surface artery and up to 4 × wider in iNPH subjects^28^, which would correspond to substantially larger PVS area ratios (3 or higher). To reflect the human scale, we here represent each PVS segment as an annular cylinder with inner radius *R*_1_ and outer radius *R*_2_ of width and area proportional to that of the corresponding blood vessel (*R*_2_ = 2*R*_1_ at baseline, *R*_2_ = 3*R*_1_ for enlarged PVS). The hydraulic resistance of annular cross-sections is 1− 6×^136^ larger than more elongated cross-sections and thus our estimates of the pressure-induced PVS velocities are conservative.

